# Chromatin mapping and single-cell immune profiling define the temporal dynamics of ibrutinib drug response in chronic lymphocytic leukemia

**DOI:** 10.1101/597005

**Authors:** André F. Rendeiro, Thomas Krausgruber, Nikolaus Fortelny, Fangwen Zhao, Thomas Penz, Matthias Farlik, Linda C. Schuster, Amelie Nemc, Szabolcs Tasnády, Marienn Réti, Zoltán Mátrai, Donat Alpar, Csaba Bödör, Christian Schmidl, Christoph Bock

## Abstract

Chronic lymphocytic leukemia (CLL) is a genetically, epigenetically, and clinically heterogeneous disease. Despite this heterogeneity, the Bruton tyrosine kinase (BTK) inhibitor ibrutinib provides effective treatment for the vast majority of CLL patients. To define the underlining regulatory program, we analyzed high-resolution time courses of ibrutinib treatment in closely monitored patients, combining cellular phenotyping (flow cytometry), single-cell transcriptome profiling (scRNA-seq), and chromatin mapping (ATAC-seq). We identified a consistent regulatory program shared across all patients, which was further validated by an independent CLL cohort. In CLL cells, this program starts with a sharp decrease of NF-κB binding, followed by reduced regulatory activity of lineage-defining transcription factors (including PAX5 and IRF4) and erosion of CLL cell identity, finally leading to the acquisition of a quiescence-like gene signature which was shared across several immune cell types. Nevertheless, we observed patient-to-patient variation in the speed of its execution, which we exploited to predict patient-specific dynamics in the response to ibrutinib based on pre-treatment samples. In aggregate, our study describes the cellular, molecular, and regulatory effects of therapeutic B cell receptor inhibition in CLL at high temporal resolution, and it establishes a broadly applicable method for epigenome/transcriptome-based treatment monitoring.

## Introduction

Chronic lymphocytic leukemia (CLL) is among the most frequent blood cancers^1^. It is characterized by clonal proliferation and accumulation of malignant B lymphocytes in the blood, bone marrow, spleen, and lymph nodes. On a cellular level, this process is driven by constitutively activated B cell receptor (BCR) signaling, which can be caused by erroneous (auto)antigen recognition and/or cell-autonomous mechanisms^2^. CLL shows remarkable clinical heterogeneity, with some patients pursuing an indolent course, while others progress rapidly and require early treatment. Extensive heterogeneity exists also at the genetic, epigenetic, and transcriptional level and has led to the identification of genetically defined CLL subtypes^3-6^ and patient-specific transcriptional programs^7-9^. Moreover, characteristic DNA methylation patterns appear to reflect differences in the CLL’s cell-of-origin^10-12^, and chromatin profiles predict the BCR immunoglobulin heavy-chain variable (IGHV) gene mutation status^13^.

Despite widespread clinical and molecular heterogeneity, therapeutic inhibition of BCR signaling has shown remarkable efficacy for CLL therapy in essentially all patients, with low rates of primary and secondary resistance. Most notably, treatment with the Bruton tyrosine kinase (BTK) inhibitor ibrutinib^14^ achieves high clinical response rates even in patients carrying genetic markers predictive of fast disease progression such as TP53 aberrations^15,16.^ As the result, ibrutinib is becoming the standard of care for a large percentage of patients with high-risk CLL.

The mechanism of action of ibrutinib is rooted in its inhibition of BTK, which leads to downregulation of BCR signaling. Previous studies have investigated specific aspects of the molecular response to ibrutinib, for example investigating immunosuppressive mechanisms^17^ and identifying decreased NF-κB signaling as a cause of reduced cellular proliferation^18-20^; but they did not map the genome-scale, time-resolved regulatory response to ibrutinib in primary patient samples. A detailed understanding of these temporal dynamics is particularly relevant given that successful ibrutinib therapy often induces an initial increase (rather than decrease) of CLL cells in peripheral blood, which can take months to resolve^21,22^. This observation has been explained by the drug’s effect on cell-cell contacts^23,24^, which triggers relocation of CLL cells from a protective microenvironment to the peripheral blood. The fact that ibrutinib induces lymphocytosis also contributes to the low correlation between the CLL cell count in the blood and the clinical response to ibrutinib therapy^22^, and there is an unmet need for early molecular markers of response to ibrutinib therapy.

To dissect the precise cellular and molecular changes induced by ibrutinib therapy, and to identify candidate molecular markers of therapy response, we followed individual CLL patients (n = 7) at high temporal resolution (eight time points) over a standardized 240-day time course of ibrutinib treatment. Peripheral blood samples were analyzed for cell composition by flow cytometry, for epigenetic/regulatory cell state by ATAC-seq^25^ on six different FACS-purified immune cell populations (158 ATAC-seq profiles in total), and for cell type specific transcriptional changes by single-cell RNA-seq^26^ applied to a subset of time points (>43,000 single-cell transcriptomes in total).

Integrative bioinformatic analysis of the resulting dataset identified a consistent regulatory program of ibrutinib-induced changes that was shared across all patients: Within the first days after the start of ibrutinib treatment, CLL cells displayed reduced NF-κB binding, followed by reduced activity of lineage-defining transcription factors, and erosion of CLL cell identity. Finally, after an extended period of ibrutinib treatment, a quiescence-like gene signature was acquired by CLL cells – and unexpectedly also by CD8^+^ T cells and other immune cell populations. This drug-induced regulatory program was present in all patients, and we were able to validate it in an independent CLL cohort. However, we observed substantial patient-to-patient variation in the speed with which these events unfold. Taking advantage of our time series data, we identified patient-specific predictors of the time to acquire an ibrutinib-induced molecular response, and we found predictive regulatory patterns already in pre-treatment samples.

In aggregate, our study provides a comprehensive, time-resolved analysis of the molecular and cellular dynamics upon ibrutinib treatment in CLL. It constitutes one of the first high-resolution, multi-omics time series of the molecular response to targeted therapy in cancer patients. The study also establishes a broadly applicable approach for analyzing drug-induced regulatory programs and identifying molecular response markers for targeted therapy. Importantly, the study’s high temporal resolution and its use of three complementary assays provided robust and informative results based on a small number of samples. The presented approach may be particularly relevant for obtaining maximum insight from early-stage clinical trials and off-label drug use involving few individual patients.

## Results

### Ibrutinib therapy induces global changes in immune cell composition and single-cell transcription profiles

To investigate the cellular dynamics and regulatory program induced by the inhibition of BCR signaling in CLL patients, we followed seven individuals from the start of ibrutinib therapy over a standardized time course of 240 days (**Figure 1a**). All patients received the same treatment regimen with daily doses of ibrutinib and underwent extensive clinical monitoring. The patients covered a range of different demographic, clinical, and genetic parameters, representative of the spectrum of refractory CLL encountered in clinical practice (**Supplementary Table 1**).

**Figure 1:**
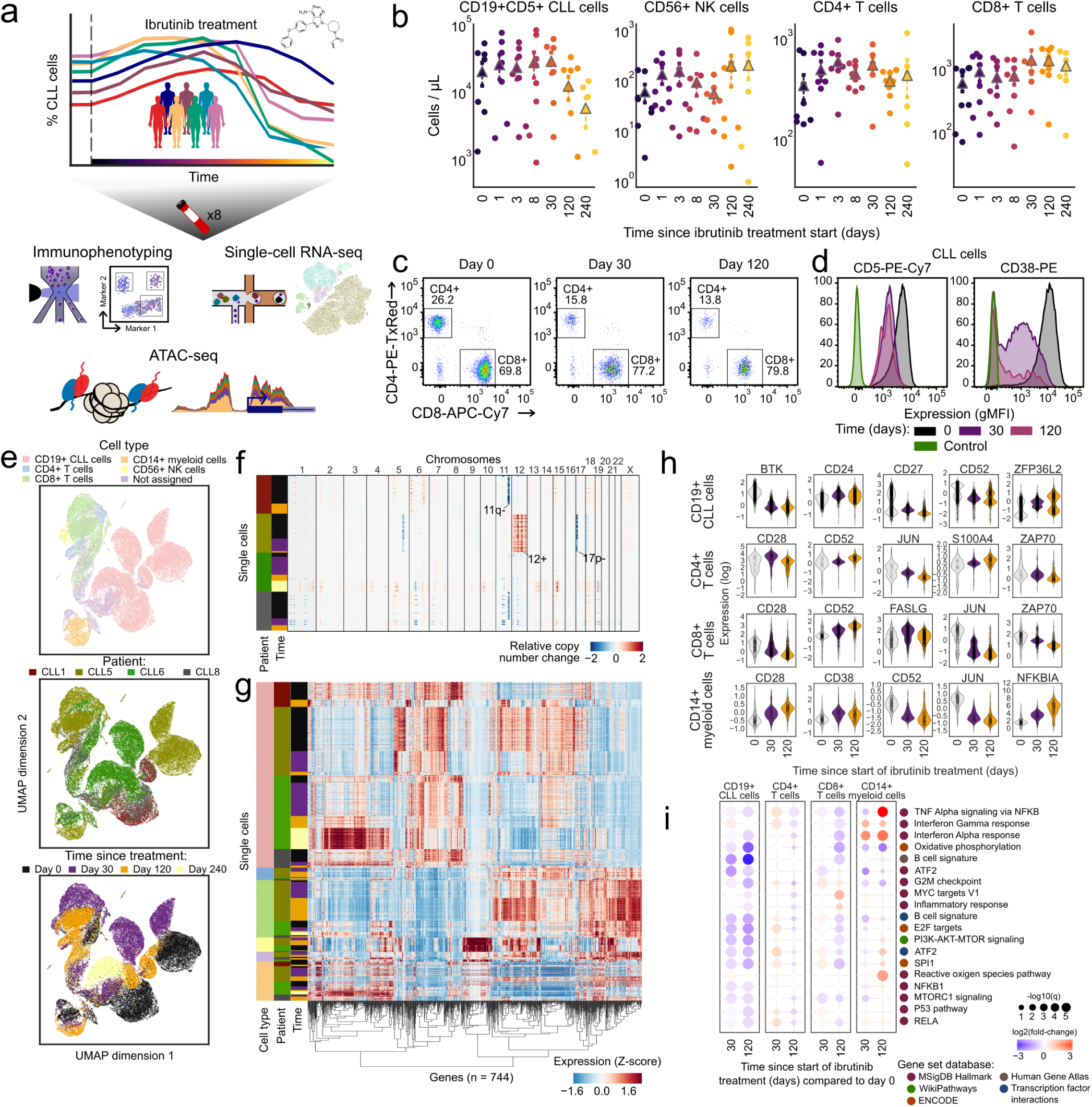
Time series analysis of the cellular and transcriptional response to ibrutinib in CLL patients identifies widespread changes in several immune cell types. **a)** Schematic representation of the study design. Peripheral blood from CLL patients undergoing single-agent ibrutinib therapy was collected at defined time points and assayed by flow cytometry (immunophenotype), single-cell RNA-seq (gene expression), and ATAC-seq (chromatin regulation). **b)** Cell type abundance over the ibrutinib time course, measured by flow cytometry. Triangles represent the mean for each time point. **c)** Flow cytometry scatterplots showing the abundance of T cell subsets for one representative patient at three time points (day 0: before the initiation of ibrutinib therapy, day 30 (120): 30 (120) days after the initiation of ibrutinib therapy). Cells positive for CD3 or CD8 were gated as indicated by the black rectangles and quantified as percentages of live PBMCs. **d)** Flow cytometry histograms showing CD5 and CD38 expression on CLL cells (pre-gated for live, single CD19^+^CD5^+^ cells) for a representative patient and three time points. **e)** Two-dimensional similarity map (UMAP projection) showing all single-cell transcriptome profiles that passed quality control. Cells are color-coded according to their assigned cell types based on the expression of known marker genes. **f)** DNA copy number profiles for CLL cells inferred from single-cell RNA-seq data, which detect three genetic aberrations common in CLL (annotated in the pane). For illustration, 2,500 randomly selected CLL cells are shown for each patient. **g)** Clustered single-cell transcriptome heatmap for the most differentially expressed genes between time points. For illustration, 20,000 cells are shown. **h)** Violin plots showing the distribution of gene expression levels for selected differentially expressed genes over the time course. **i)** Differential gene expression signatures in four cell types, comparing each sample to the matched pre-treatment sample and averaging across patients. Patient-individual data are shown in Supplementary Figure 6.

For all patients and up to eight time points (0, 1, 2, 3, 8, 30, 120/150, 240 days after the start of ibrutinib therapy), we performed immunophenotyping by flow cytometric analysis of peripheral blood mononuclear cells (PBMCs), systematically quantifying changes in cell composition in response to ibrutinib therapy (**Supplementary Figure 1a and Supplementary Table 2**). A gradual decrease in the percentage of CLL cells was observed over time (**Figure 1b**), but with extensive temporal heterogeneity across patients (**Supplementary Figure 1b,c**). The progressive reduction in the percentage of CLL cells coincided with an increase in the percentage of non-malignant natural killer (NK) and T cell populations, consistent with a recent report^23^. This trend was most visible for CD8^+^ T cells (**Figure 1b,c and Supplementary Table 2**), while CD4^+^ T cells remained largely unaffected. Although these differences were not statistically significant due to small cohort size, they were consistent with published data and thus provided validation and cellular characterization of our cohort and time course of ibrutinib therapy.

Based on flow cytometry, we also observed a statistically significant loss of CLL-associated surface receptors (CD5, CD38), which was specific to CLL cells (**Figure 1d**, **Supplementary Figure 2**, and **Supplementary Table 3**). To investigate the ibrutinib-induced changes in gene expression more systematically – and simultaneously in CLL cells as well as in matched non-malignant immune cells, we performed droplet-based single-cell RNA-seq^26^ on the total PBMC population for a subset of patients and time points (**Supplementary Table 4**). Overall, ∼43,000 single-cell transcriptomes passed quality control (**Supplementary Figure 3a,b**) and were integrated into a two-dimensional map using the UMAP method for unsupervised dimensionality reduction (**Figure 1e**).

Cell type specific marker genes (e.g., CD79A, CD3D, CD14, and NKG7) were readily detectable in the single-cell RNA-seq data and were largely unaffected by ibrutinib treatment (**Supplementary Figure 3c**), thus allowing for robust marker-based assignment of cell types. Cell counts inferred from single-cell RNA-seq were almost perfectly correlated with those obtained by flow cytometry (Spearman’s ρ = 0.95, **Supplementary Figure 3d**), which provided independent validation of our single-cell RNA-seq dataset. We were also able to infer patient-specific copy number aberrations from the single-cell RNA-seq data (**Figure 1f**), which identified characteristic CLL-specific chromosomal aberrations including the deletion of chromosome 11q and 17p, and trisomy of chromosome 12.

Comparing the single-cell transcriptomes for each sample and cell type to the patient’s corresponding pre-treatment (day 0) sample (**Supplementary Figure 3e-j and Supplementary Table 5**), we found cell type specific trends in the molecular response to ibrutinib therapy (**Figure 1g-h and Supplementary Figure 4**). In CLL cells, we observed reduced expression of the ibrutinib target BTK, of CD52 (a CLL disease activity marker^27^), and of CD27 (a regulator of B cell activation^28^). Among the non-malignant immune cell types, CD8^+^ T cells were most strongly affected, which included downregulation of genes important for immune cell activation such as CD28, JUN, and ZAP70. This pattern was shared to a lesser extent by CD4^+^ T cells, while CD14^+^ cells were characterized by strong upregulation of the NF-KB regulator NFKBIA.

Looking beyond individual genes, we further characterized the molecular response to ibrutinib by quantifying the transcriptome dynamics of predefined gene sets and transcriptional modules relevant to CLL and immunity (**Figure 1i and Supplementary Figure 5, 6**). We observed robust downregulation of B cell specific genes in CLL cells, including target gene sets of NF-κB subunits RELA and NF-κB1, as well as target gene sets of the well-established NF-κB associated transcription factors ATF2 and SPI1/PU.1. Genes involved in oxidative phosphorylation were also downregulated, consistent with widespread dampening of cellular activities in CLL cells under ibrutinib therapy. Among the non-malignant immune cell types, CD8^+^ T cells showed broad downregulation that was less pronounced but similar to the response observed in CLL cells, and CD14^+^ monocytes/macrophages showed specific upregulation of inflammatory response signatures including interferon gamma, TNF, and NF-κB signaling.

In summary, immunophenotyping and single-cell RNA sequencing over a dense time course of ibrutinib therapy uncovered widespread changes not only in CLL cells, but also in non-malignant immune cells. Most notably, we observed downregulation of NF-κB signaling and loss of B-cell surface markers in CLL, suggesting these are key contributors to the progressive reduction of the CLL cell fraction over time, and we observed a surprising degree of transcriptional change in non-CLL immune cells concomitant with an increase in the CD8^+^ T cell fraction.

### Chromatin mapping in CLL cells defines an ibrutinib-induced regulatory program leading to loss of B cell identity

To dissect the regulatory basis of the ibrutinib-induced changes in the CLL cell transcriptomes and immunophenotypes, we performed ATAC-seq on the FACS-purified CD19^+^CD5^+^ cell compartment over the ibrutinib time course (**Figure 2a, Supplementary Figure 7, and Supplementary Table 6**). We modeled the temporal progression as Gaussian processes (a statistical method for handling time series data^29^) and identified 6,797 genomic regions that underwent significant changes in chromatin accessibility in response to ibrutinib treatment (**Supplementary Table 7**). Four major clusters were detected among these genomic regions (**Figure 2b)**: (i) regions that gradually lost chromatin accessibility (n = 3,412); (ii) regions that gradually gained chromatin accessibility (n = 2,199); (iii) regions that followed a bimodal, oscillating pattern (n = 369); and (iv) regions characterized by a peak in chromatin accessibility around 30 days after the start of ibrutinib treatment (n = 354).

**Figure 2:**
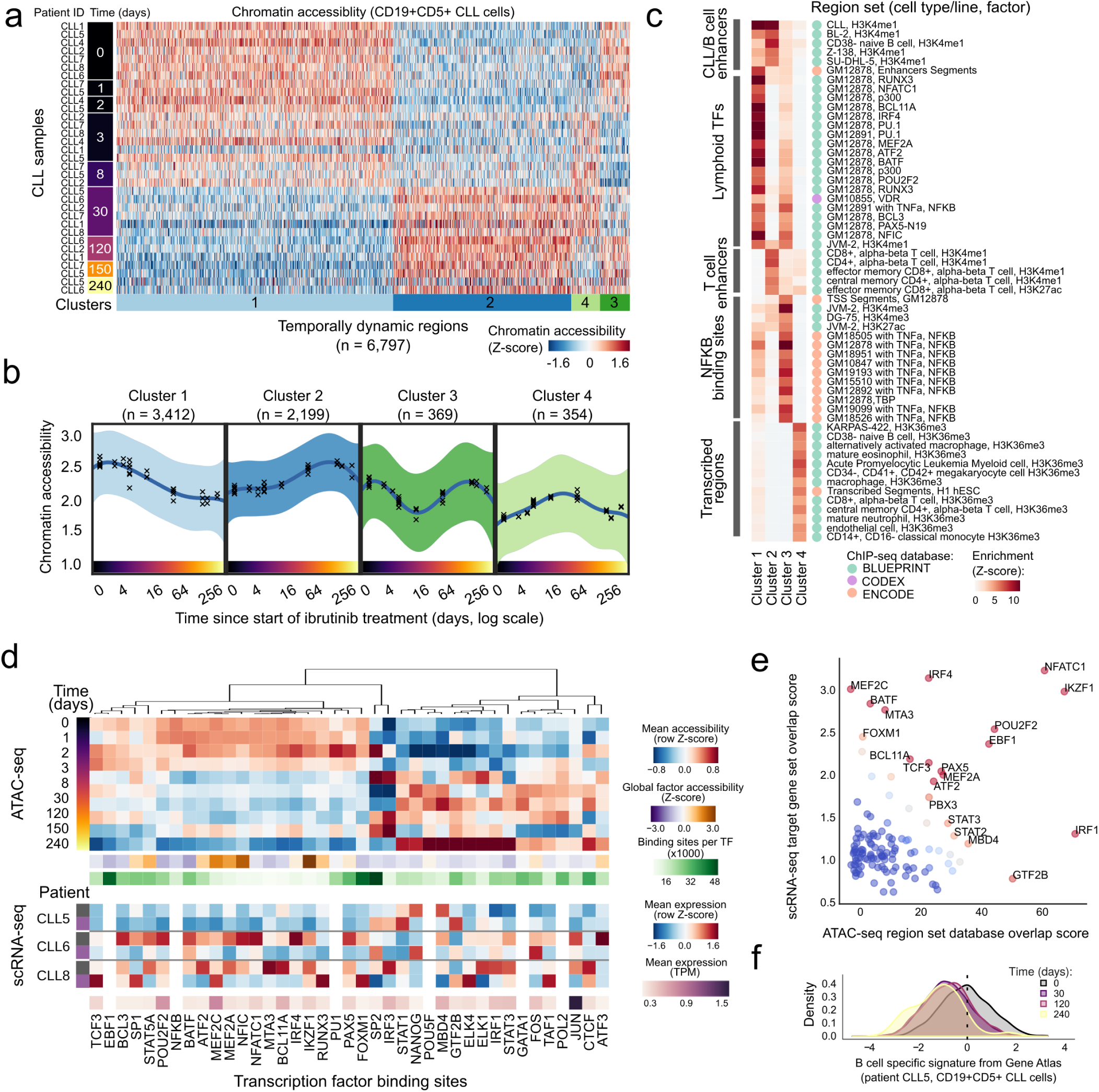
Integrated analysis of chromatin accessibility and gene expression in CLL cells uncovers a consistent regulatory program induced by ibrutinib therapy. **(a)** Heatmap showing changes in chromatin accessibility for CLL cells over the time course of ibrutinib treatment. **(b)** Mean chromatin accessibility across patients plotted over the ibrutinib time course in dynamically changing regulatory regions, highlighting the non-linear aspect of the ibrutinib effect on the chromatin. Crosses represent samples from a single patient at a specific time point, and 95% confidence intervals are shown as colored shapes. **(c)** Region set enrichments for the clusters of dynamic regions, calculated using the LOLA software. Enrichment p-values were Z-score transformed per column. **d)** Heatmaps showing mean chromatin accessibility of regulatory regions overlapping with putative binding sites, expression of the corresponding transcription factor, and total number of its binding sites. Clustering was performed on the mean chromatin accessibility values. **e)** Scatterplot showing differential regulation of transcription factors upon ibrutinib treatment. The x-axis displays the enrichment of transcription factors enriched in the LOLA analysis, and the y-axis displays the enrichment of their target genes among the differentially expressed genes. **f)** Gene expression histogram across CLL cells in one patient, demonstrating the decline of a B cell-specific expression signature over the time course of ibrutinib treatment. For illustration, the patient with most time points in the single-cell RNA-seq analysis (CLL5) is displayed.

We inferred the putative regulatory roles of these four clusters by region set enrichment analysis using the LOLA software^30^. LOLA identified those region sets out of a large reference dataset that showed significant overlap with the regions of the respective cluster (**Figure 2c**). Cluster 1 (decrease in chromatin accessibility) was strongly enriched for binding sites of transcription factors with a role in lymphoid differentiation and gene regulation, and also for enhancers specific to CLL cells and/or B cells. Cluster 2 (increase in chromatin accessibility) was enriched for B cell as well as T cell specific enhancers. Cluster 3 (bimodal, oscillating chromatin accessibility) was enriched for NF-κB binding sites. Lastly, Cluster 4 (peak in chromatin accessibility around day 30) was enriched for intragenic, transcribed regions marked by histone H3K36me3 in a range of hematopoietic cell types.

To identify potential regulators of the ibrutinib-induced modulation of CLL cell state, we focused on the enriched transcription factors (from **Figure 2c**) and estimated their dynamic changes in global binding activity over the ibrutinib time course, aggregating the ATAC-seq signal across each factor’s putative binding sites (as defined by publicly available ChIP-seq data). As expected, several key transcription factors involved in B cell development (including NF-κB and PAX5) and B cell proliferation (including MEF2C and FOXM1) showed marked reduction of chromatin accessibility at their binding sites (**Figure 2d and Supplementary Figure 8a**). This effect was shared by CLL cells and non-malignant B cells but not observed in other immune cell types.

Integrative analysis of chromatin accessibility and cell type specific transcription further refined this picture. When we performed parallel enrichment analysis for transcription factors and their putative binding sites (**Figure 2e**), we observed concerted changes for key regulators of B cell development such as BCL11A, EBF1, IKZF1, IRF4, MEF2A, NFATC1, PAX5, and POU2F2, indicating that BTK inhibition may trigger loss of B cell identity in CLL cells. In support of this interpretation, we found global B cell specific gene expression signatures consistently downregulated upon ibrutinib treatment in CLL cells (**Figure 2f and Supplementary Figure 8b**).

Taken together, these results define a characteristic temporal order in which the ibrutinib-induced changes in CLL cells unfold. Already after one day of ibrutinib treatment, CLL cells showed a reduction in chromatin accessibility at NF-κB binding sites. This was followed by a gradual decrease in chromatin accessibility at binding sites of transcription factors that are regulated by NF-κB (PU.1^31^, IRF4^32,33^) or interact with NF-κB (ATF2^34^). Moreover, we observed widespread reduction in B cell specific regulatory activity including decreased chromatin accessibility at B cell specific elements and at the binding sites of B cell transcription factors such BCL11A, NFATC1, and RUNX3, which was also reflected at the transcriptional level. These results highlight NF-κB mediated loss of B cell identity as the central cell-intrinsic change in CLL cells from patients under ibrutinib therapy.

### Analysis of five immune cell types identifies ibrutinib-induced acquisition of a shared quiescence gene signature

To characterize the effect of ibrutinib therapy on gene regulation in non-malignant immune cells, we performed ATAC-seq on FACS-purified CD19^+^CD5^-^ B cells, CD3^+^CD4^+^ T helper cells, CD3^+^CD8^+^ cytotoxic T cells, CD56^+^ NK cells, and CD14^+^ monocytes/macrophages from the same patients and time points (**Supplementary Table 8**). We identified a total of 12,574 temporally dynamic regulatory regions in these five cell types (**Figure 3a,b; Supplementary Figure 9**; **Supplementary Table 9**). Unsupervised clustering detected shared temporal dynamics across these cell types, with sets of regions showing gradually decreased or increased chromatin accessibility over the time course, and a bimodal, wave-like cluster that was characterized by an initial decrease followed by a subsequent increase in chromatin accessibility (**Figure 3c and Supplementary Figure 9a-c**). Despite the shared temporal dynamics, the affected regions were highly cell type specific (**Supplementary Figure 9d**), suggesting that the different immune cell types react in characteristic ways to the direct and indirect effects of ibrutinib treatment.

**Figure 3:**
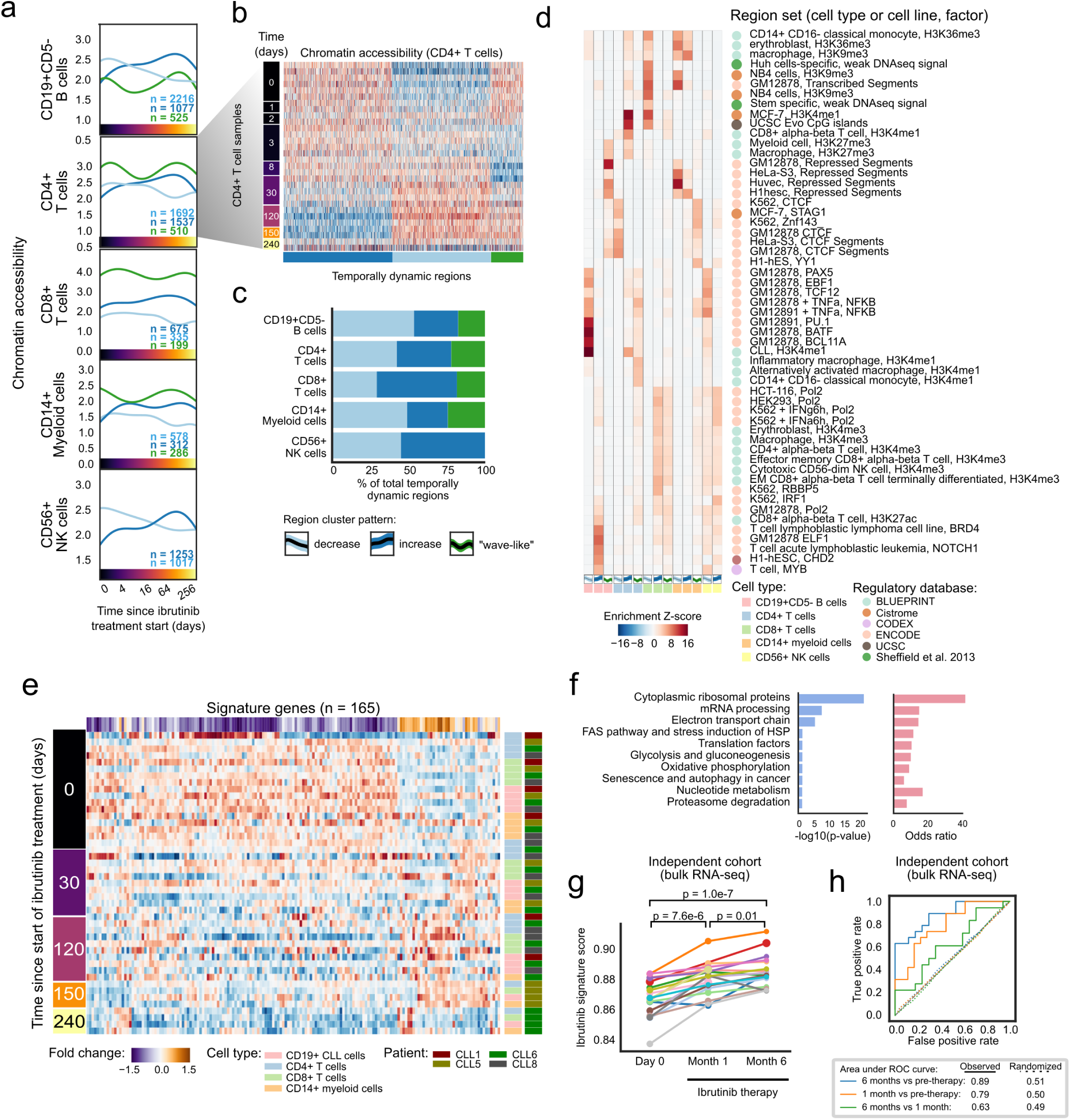
Integrated analysis of chromatin accessibility and gene expression for five immune cell types identifies regulatory changes in response to ibrutinib that converge on a shared quiescence-like gene signature. **a)** Mean chromatin accessibility across patients plotted over the ibrutinib time course for clusters of dynamically changing regulatory regions in five immune cell types. **b)** Heatmap of chromatin accessibility of CD4+ cells, illustrating dynamic regulation over the ibrutinib time course. **c)** Stacked bar plots indicating the percentage of dynamically changing regions in each cluster. **d)** Region set enrichments for the clusters of dynamically changing regions, calculated using the LOLA software and publicly available region sets as reference (mainly based on ChIP-seq data). Enrichment p-values were Z-score transformed per column. **e)** Heatmap showing mean expression levels for genes that were differentially expressed over the ibrutinib time course when combining the data for CLL cells and for the five non-malignant immune cell types. Values represent column Z-scores of gene expression. **f)** Gene set enrichments for genes downregulated across cell types, using WikiPathways as reference (Fisher’s exact test, left: FDR-corrected p-value, right: odds ratio as a measure of effect size). **g)** Expression score for the gene signature (as shown in panel e) in an independent cohort, calculated from bulk RNA-seq data for PBMCs collected before the start of ibrutinib therapy and at two subsequent time points. Significance was assessed using a paired t-test. **h)** ROC curve illustrating the prediction performance of the gene signature (from panel e) for classifying samples in the independent validation cohort into those collected before ibrutinib treatment and those collected during ibrutinib treatment. As negative controls, the prediction was repeated 100 times with permuted class labels for each combination of time points, and the mean ROC curves across iterations are shown as dotted lines.

Of the five non-malignant immune cell types, CD19^+^CD5^-^ B cells were most strongly affected by ibrutinib therapy, consistent with the major role of BCR signaling and the ibrutinib target BTK that non-malignant B cells share with CD19^+^CD5^+^ CLL cells. Regions with decreasing chromatin accessibility in the non-malignant B cells were enriched for the same set of transcription factor binding sites as the CLL cells, and also, to a lesser extent, for NF-κB binding sites (**Figure 3d**). In contrast, we detected fewer regions with increasing chromatin accessibility upon ibrutinib treatment, and those regions lacked distinctive patterns of functional enrichment, suggesting that they are indirect effects downstream of the cells’ direct response to ibrutinib treatment (**Supplementary Table 9**).

Biologically interesting changes were not restricted to B cells. For example, regions with decreasing chromatin accessibility in CD4^+^ T cells upon ibrutinib treatment were enriched for binding sites of CTCF and RAD2, which are involved in three-dimensional chromatin organization; and regions with decreasing chromatin accessibility in CD8^+^ T cells were enriched for histone marks associated with repressed chromatin in other cell types (**Figure 3d**). Conversely, regions with increased chromatin accessibility in CD4^+^ T cells over the ibrutinib time course were enriched for interferon signaling and open, promoter-associated chromatin in T cells, while the enrichment observed for CD8^+^ T cells included CpG islands and H3K4me1-marked regulatory regions (**Figure 3d**).

Despite the cell type specific effects of ibrutinib therapy on CLL cells and non-malignant immune cells, we also identified a characteristic set of genes that underwent consistent changes across all of the investigated cell types (**Figure 3e and Supplementary Figure 9e**). This shared ibrutinib response signature was enriched for genes involved in ribosomal functions, mRNA processing, oxidative phosphorylation/metabolism, translation factors, senescence, and autophagy (**Figure 3f**). For example, the shared ibrutinib response signature included CD44, a pan-lymphocyte cell adhesion molecule; CD99, a regulator of leukocyte migration, T cell adhesion, and cell death; CD37, which mediates the interaction of B and T cells; various surface proteins involved in cell adhesion (CD52, CD164, ICAM3, ITGB7); the protein tyrosine kinase FGR, which is a negative regulator of cell migration; TPT1 (a regulator of cellular growth and proliferation); and several factors involved in protein translation (EEF2, EID1, EIF1, EIF3E) as well as ribosomal proteins (**Supplementary Figure 10a**).

Interestingly, we also found genes involved in senescence and/or quiescence, namely CXCR4, a chemokine receptor required for hematopoietic stem cell quiescence^35,36^; ZFP36L2, an RNA binding protein that promotes quiescence in developing B cells^37^; and HMGB2, a chromatin protein involved in the regulation of gene expression in senescent cells^38^ (**Supplementary Figure 10**). Together, these results suggest that CLL cells and non-malignant immune cells respond to ibrutinib therapy with shared transcriptional changes that include downregulation of genes involved in leukocyte function and cell-cell interactions, as well as upregulation of genes involved in quiescence and cellular senescence.

To assess the reproducibility of this shared ibrutinib response signature in an independent validation cohort, we utilized recently published bulk RNA-seq data for PBMCs from CLL patients (n = 19) that underwent single-agent ibrutinib treatment at a different medical center^20^. We indeed observed consistent changes in the expression of our gene signature for the vast majority of patients from the validation cohort (**Figure 3g**). The difference was statistically significant at both time points compared to day 0 (month 1: p = 7.6e-6; month 6: p = 1.0e-7; paired *t*-test), and an accurate distinction was possible between patient samples collected before and during ibrutinib therapy (receiver operating characteristic area under curve values of 0.89 and 0.79, respectively) (**Figure 3h**).

In summary, our data show that ibrutinib therapy induces time-dependent regulatory changes not only in CLL cells but also in other immune cell types. Changes in non-malignant B cells mirrored those in CLL cells (albeit with a weaker NF-κB signature), while CD4^+^ T cells, CD8^+^ T cells, NK cells, and myeloid cells responded in cell type specific ways. We further identified and validated a gene expression signature that captures broad ibrutinib-induced downregulation of immune cell functions and acquisition a quiescent-like state in response to ibrutinib therapy.

### Prediction of patient-to-patient variability over the time course provides a molecular marker of ibrutinib response

Our data and analyses strongly support the existence of a consistent, ibrutinib-induced regulatory program that is shared across all patients. Nevertheless, we also observed substantial patient-to-patient variability at the genetic (**Figure 1f)**, transcriptional (**Figure 3g**), chromatin-regulatory (**Figure 2d**), and cellular level (**Supplementary Figure 1b**). This heterogeneity in the presence of a shared regulatory program could be explained by patient-to-patient differences in the speed of progression along the regulatory program. Analyzing the molecular progression may therefore provide us with an opportunity to monitor or even predict, based on molecular profiles, which patients pursue a faster or slower time course toward a sustained cellular response under ibrutinib therapy.

Along these lines, we first investigated whether there were changes in the subclonal composition of CLL cells under ibrutinib treatment. To that end, we analyzed the copy number profiles inferred from the single-cell RNA-seq data and indeed observed subclonal genetic differences within patients over the time course (**Supplementary Figure 10a-d**). We also inferred the molecular response to ibrutinib therapy for each individual CLL cell, based on the expression intensity of our validated ibrutinib response signature (**Figure 3e**) in the single-cell transcriptomes. When we correlated this “ibrutinib molecular response score” with the single-cell copy number profiles, we did not observe a clear association between individual copy number aberrations and the strength of the molecular response to ibrutinib in single cells (**Supplementary Figure 10e-h**). However, we did observe an increase of subclonal genetic diversity of the time course of ibrutinib response, based on a quantitative measure that we validated on simulated data and on the changing ratio of CLL cells versus non-malignant cells in our time course (**Figure 4a and Supplementary Figure 10i-l)**. This change of subclonal genetic heterogeneity within patients was indeed positively associated with a strong cellular response to ibrutinib treatment as measured the flow cytometry data (**Figure 4b**).

**Figure 4:**
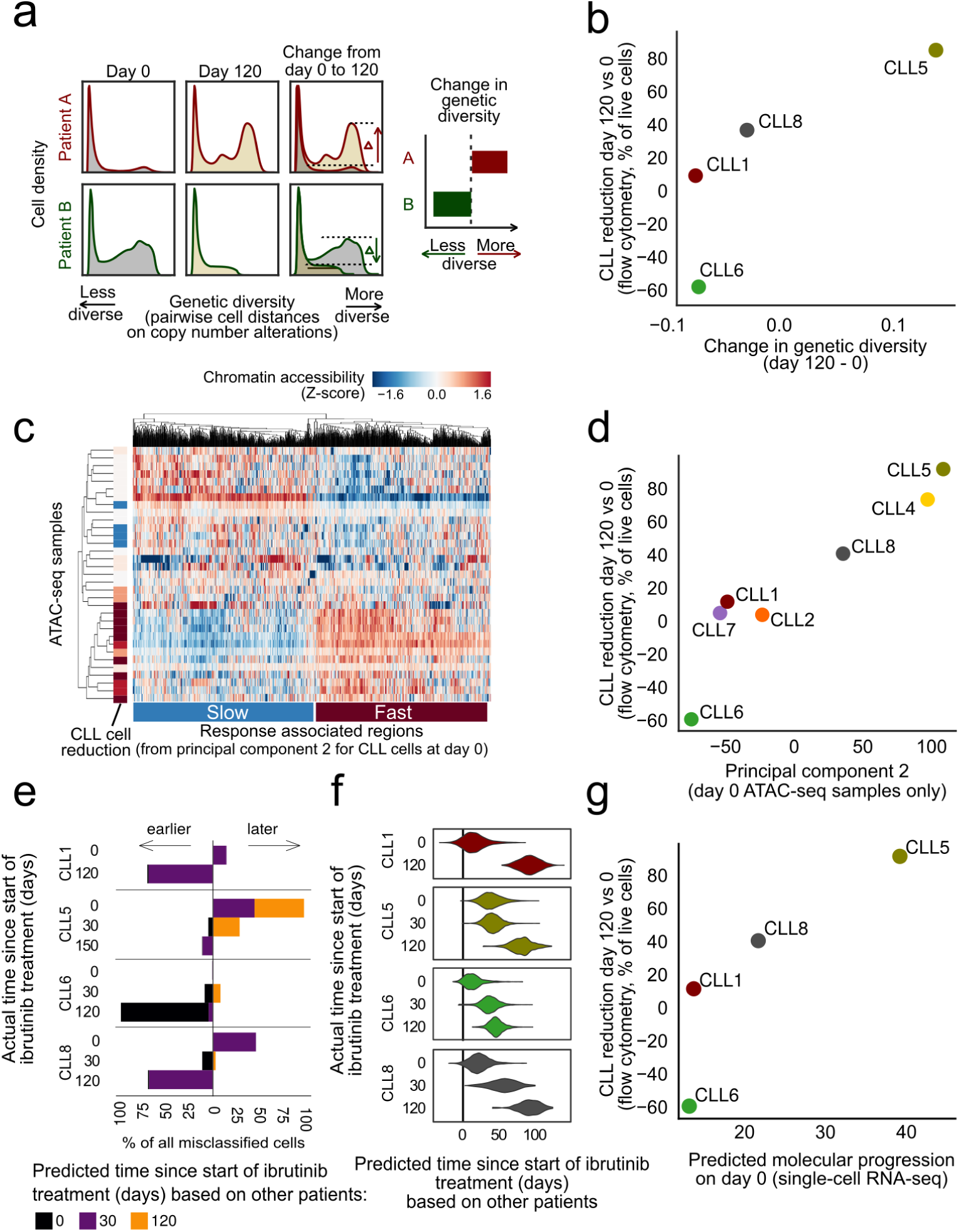
Analysis of copy number profiles, chromatin accessibility, and single-cell transcriptomes identified patient-specific associations with the speed of the cellular response to ibrutinib therapy. **a)** Computational approach to quantify changes in genetic diversity based on copy number profiles inferred from the single-cell RNA-seq data. Shifts in the distribution of pairwise distance similarities between time points indicate changes in the genetic diversity of the cell population. **b)** Scatterplot comparing across patients the change in genetic diversity between day 0 and 120/150 of ibrutinib treatment (x-axis) with the change in the CLL cell percentage on day 120 /150 of ibrutinib treatment compared to day 0 as measured by flow cytometry (y-axis). **c)** Clustered heatmap showing chromatin accessibility profiles for CLL cells at day 0 for the top 1000 genomic regions associated with the second principal component for these profiles (from **Supplementary Figure 11a**), annotated on the left with the change in CLL cell fraction (as in panel b). **d)** Scatterplot comparing across patients the average chromatin accessibility across regions linked to the second principal component (as in panel c, x-axis) with the change in CLL cell fraction (as in panel b, y-axis). **e)** Stacked bar charts showing the number and direction of deviations from the actual collection time point when predicting time points in each patient after training the classifier in all other patients. **f**) Violin plots showing the predicted (x-axis) and actual (y-axis) number of days under ibrutinib therapy in each patient. Predictions are derived from regression models trained on all other patients. **g)** Scatterplot comparing the predicted time under ibrutinib therapy (from panel f, x-axis) with the change in CLL cell fraction (as in panel b, y-axis).

Second, we investigated the association of chromatin accessibility in CLL cells at day 0 with a range of patient-specific characteristics. To that end, we performed principal component analysis on the chromatin profiles for all patients and cell types, and we tested for statistical associations with the clinical annotation data (**Supplementary Figure 11a**). We observed a strong association between the second principal component of the chromatin profiles in CLL cells at day 0 and the cellular response to ibrutinib treatment at day 120, suggesting that this chromatin signature provides an epigenomic marker for the subsequent cellular response to ibrutinib treatment (**Figure 4c,d**). This chromatin signature separated patients into fast versus slow responders to ibrutinib treatment independently of other clinical annotations (**Supplementary Figure 11b**). Genomic regions associated with a slow response to ibrutinib treatment showed essentially the same enrichment as those that were downregulated in CLL cells (**Figure 2c**), including preferential overlap with a broadly active state of cellular activity (**Supplementary Figure 11c)**.

Third, we employed our single-cell RNA-seq dataset to derive and evaluate gene expression signatures that capture the molecular response to ibrutinib treatment in individual cells. Using machine learning, we predicted the time of sample collection (day 0, 30, or 120/150) for each of the ∼19,000 single-cell transcriptomes for CLL cells from four donors. Both support vector machines and elastic net classifiers achieved excellent prediction performance with cross-validated test-set ROC area under curve (AUC) values in the range of 0.975 to 0.999, and these results were robust to differences in the number of detected genes among the single-cell transcriptome profiles (**Supplementary Figure 12a**). Our observations indicate that the transcriptome profiles of single CLL cells undergo changes that precisely reflect the duration of ibrutinib therapy – which we can exploit for molecular staging of the patient-specific ibrutinib response. Using a classifier that was trained and evaluated by patient-stratified cross-validation, we observed that cells from specific patients were consistently predicted to have progressed faster (CLL5) or slower (CLL6) along the trajectory of the transcriptional response to ibrutinib treatment (**Figure 4e**), indicating that individual patients indeed follow their own timelines in the molecular response to ibrutinib therapy.

Finally, for a more quantitative assessment of these temporal dynamics, we trained and evaluated regression models that predict the precise time (i.e., number of days) after the start of ibrutinib therapy for each individual CLL cell transcriptome. We observed excellent test set prediction performance for three patients (CLL1, CLL6, and CLL8), with r^2^ values (i.e., percent variance explained) of 92.3%, 84.2%, and 78.1%, respectively (**Supplementary Figure 12b-c**). Lower performance was observed for CLL5 (r^2^ = 36.6%), where the day-0 time point already showed a signature reminiscent of ibrutinib treatment (**Figure 4f)**. Consistent with the results of the classification analysis (**Figure 4e**), the regression models predicted individual patients progressing faster (CLL5) or slower (CLL6) along the trajectory of transcriptional response, while the two remaining samples (CLL1, CLL8) followed similar timelines (**Figure 4f**). When we compared predictions based on CLL cell transcriptomes at day 0 across patients, we found that the observed molecular signature prior to the start of ibrutinib treatment indeed anticipated the subsequent cellular response (i.e., reduction of CLL cells on day 120/150 compared to day 0) (**Figure 4g**).

These results indicate that genetic, epigenetic, and transcriptional variation between patients capture inter-individual differences in the response to ibrutinib treatment and may provide molecular markers that predict the time to a strong cellular response for individual patients.

## Discussion

Multi-omics analysis of clinical time courses provides an effective approach for dissecting the molecular response to targeted therapy, allowing us to define the temporal order of events and to unravel relevant regulatory programs. Here, we applied flow cytometry, single-cell RNA-seq, and chromatin mapping in six FACS-purified cell types to a dense time course of CLL patients starting ibrutinib therapy. These three assays provide comprehensive and complementary information comprising the cellular response (flow cytometry), transcriptional changes across all major immune cell populations (single-cell RNA-seq), and the underlying chromatin dynamics that may explain and predict the observed changes in transcription regulation and epigenetic cell state (ATAC-seq).

Integrative bioinformatic analysis identified a characteristic regulatory program that was shared across all patients. Among the earliest changes following the start of ibrutinib therapy, we observed a decrease of NF-κB binding in CLL cells, which was followed by a rapid reduction in the regulatory activity of transcription factors involved in B cell development and function (such as EBF1, FOXM1, IRF4, PAX5, and PU.1). This decrease was accompanied by (and it likely caused) downregulation of CLL-specific gene signatures and a decrease in surface marker levels (including CD5 and CD19), which together indicate a broad erosion of CLL cell identity.

Ibrutinib-induced changes were not exclusive to CLL cells but shared by several other immune cell types. Non-malignant B cells largely mirrored the changes observed in CLL cells – which was expected given the role of the ibrutinib target BTK in BCR signaling. We also observed a dampening effect of ibrutinib on immune pathway regulation in CD8^+^ T cells, while there was an increase of inflammatory gene signatures in monocytes/macrophages. The changes in immune cell types that do not express BTK could be due to a combination of direct effects via ibrutinib’s promiscuous inhibition of kinases other than BTK (including BLK, BMX, ITK, TEC, TXK, and EGFR^39^) and indirect effects arising from the ibrutinib-induced relocation of CLL cells from the protective microenvironment into the peripheral blood.

Interestingly, for both CLL cells and for non-malignant immune cell populations, sustained ibrutinib therapy eventually resulted in the acquisition of a shared, quiescence-like gene signature. We validated this gene signature in an independent clinical cohort and confirmed its reproducibility. Closer inspection of the contributing genes may help explain certain cellular and clinical phenotypes observed in CLL patients under ibrutinib therapy, including changes in the immune microenvironment^40^ and increased susceptibility to infections^41-43^. For example, CD99 downregulation indicates that Fas-mediated T-cell death may be impaired^44,45^, which has been proposed as a cause of CD8^+^ T cell accumulation in peripheral blood^23^. Moreover, two genes in the signature (CXCR4 and ZNF36L2) have established biological functions in senescence and quiescence of hematopoietic stem cells and lymphocytes. Ibrutinib is known to inhibit CXCR4-mediated expression of CD20 in CLL cells^46^, which could have implications for ongoing trials combining ibrutinib and anti-CD20 antibodies in CLL (e.g., NCT02007044).

Our comprehensive, time-resolved, multi-omics analysis of ibrutinib therapy thus provides integration and context for previous studies that have focused on specific aspects of the response to ibrutinib, including reduced proliferation^19^, decreased cell-cell contacts^23,24^, and downregulation of NF-κB^18-20^. Moreover, the identification of a regulatory program that was shared across all patients allowed us to explore patient-to-patient heterogeneity in the speed with which this program is executed, suggesting that it may be feasible to define predictive molecular markers for the cellular response to ibrutinib therapy. Most notably, our chromatin analysis identified a patient-specific signature present prior to treatment that correlated with the speed of CLL clearance, and our single-cell RNA-seq data predicted the cellular response measured 120/150 days after the start of ibrutinib therapy. While these results remain exploratory due to the small number of patients in our study, they raise the future perspective of molecular response monitoring and prediction for a growing class of targeted cancer therapies that are not primarily cytotoxic and for which simple cell-based biomarkers (such as leukemic cell count or minimal residual disease) are poor predictors of the eventual clinical response.

While we consider our approach broadly applicable in the context of targeted therapies and precision oncology, the following limitations apply: First, such comprehensive profiling (up to 8 time points, dozens of genome-wide ATAC-seq profiles, and thousands of single-cell transcriptomes per patient) is currently feasible only for a relatively small number of patients. This precludes systematic interaction analysis with genetic risk markers for CLL. (However, recent studies have shown that established prognostic markers have lost much of their predictive power with ibrutinib therapy^47,48^, and the same may apply to other emerging treatments such as CAR T cell therapy^49^.) Second, while we found evidence of subclonal heterogeneity in our single-cell transcriptome data, the current throughput of single-cell RNA-seq does not (yet) enable deep characterization of the subclonal architecture. Third, most patients that start ibrutinib treatment have previously been treated with other drugs or drug combinations (our cohort: 1 to 5 prior treatments), which may explain some of the differences in the speed of the molecular response to ibrutinib in the individual patients. Fourth, time series data supports only a weak form of causal inference (Granger causality^50,51^), where earlier events may cause later events but not vice versa (e.g., the observed decrease in NF-κB binding was followed by a downregulation of B cell transcription factors and an erosion of B cell identity among the CLL cells). The results should therefore be considered causal in a strict biological sense only after mechanistic experimental validation in suitable disease models. With these limitations taken into account, we expect that the presented approach will readily generalize to other targeted therapies, defining shared regulatory programs and identifying molecular markers of the temporal dynamics and response to targeted therapy.

In conclusion, our study demonstrates the power of high-throughput assays combined with integrative bioinformatic analysis for dissecting the regulatory impact of targeted therapies. A strength of this approach is the level of detail and biological insight that can be obtained from a small number of patients, which makes it well-suited for applications in personalized medicine, where each patient may behave differently. Moreover, the approach appears promising for early-stage clinical trials of new targeted therapies, where it is critical to obtain a robust assessment of the induced molecular and cellular dynamics, in order to inform dose finding and to provide biomarker candidates for molecular response monitoring.

## Methods

### Sample acquisition and clinical data

All patients were treated at the Department of Haematology and Stem Cell Transplantation, Central Hospital of Southern Pest, Budapest, Hungary, according to the revised guidelines of the International Workshop Chronic Lymphocytic Leukemia/National Cancer Institute^52^. The study was approved by the ethical committees of the contributing institutions (Dél-Pesti Centrumkórház, Semmelweis University, and Medical University of Vienna). Informed consent was obtained from all participants.

### Flow cytometry and fluorescence activated cell sorting (FACS)

Patient PBMCs were thawed and washed twice with PBS containing 0.1% BSA and 5 mM EDTA (PBS + BSA + EDTA). Cells were then incubated with anti-CD16/CD32 (clone 93, Biolegend) to prevent nonspecific binding. Single-cell suspensions were stained with combinations of antibodies against CD3 (FITC, clone UCHT1), CD4 (PE-TxRed, clone OKT4), CD5 (PE-Cy7, clone UCHT2), CD8 (APC-Cy7, clone SK1), CD14 (PerCp-Cy5.5, clone M5E2), CD19 (APC, clone HIB19), CD25 (PE-Cy7, clone BC96), CD38 (PE, clone HB-7), CD45RA (PerCp-Cy5.5, clone HI100), CD45RO (AF700, clone 304218), CD56 (AF700, clone NCAM16.2), CD127 (APC, clone A019D5), CD197 (CCR7, PE, clone G043H7), and DAPI viability dye (all from Biolegend) for 30 min at 4 °C followed by two washes with PBS + BSA + EDTA. For flow cytometry, cells were acquired with an LSRFortessa Cell Analyzer (BD). For FACS, cells were sort-purified with a MoFlo Astrois (Beckman Coulter) using the gating strategy depicted in **Supplementary Figure 1a**. Data analysis was performed with the FlowJo (Tree Star) software. In Figure 1d, control cells in CD5-PE-Cy7 channel are CD14+ myeloid cells, and control cells in CD38-PE channel are CD3+CD4-CD8-cells; these are cell populations which are known to not express the respective markers and based on which the background levels can be estimated.

### Droplet-based single-cell RNA-seq

Single-cell libraries were generated using the Chromium Controller and Single Cell 3’ Library & Gel Bead Kit v2 (10x Genomics) according to the manufacturer’s protocol. Briefly, an aliquot of patient PBMCs was stained with DAPI for discrimination between live and dead cells, and a maximum of 100,000 live, doublet-excluded cells were sorted into 1.5 ml tubes. Cells were pelleted by centrifuging for 5 min at 4 °C at 300 x g and resuspended in PBS with 0.04% BSA. Up to 17,000 cells suspended in reverse transcription reagents, along with gel beads, were segregated into aqueous nanoliter-scale gel bead-in-emulsions (GEMs). The GEMs were then reverse-transcribed in a C1000 Thermal Cycler (Bio-Rad) programmed at 53 °C for 45 min, 85 °C for 5 min, and hold at 4 °C. After reverse transcription, single-cell droplets were broken, and the single-strand cDNA was isolated and cleaned with Cleanup Mix containing Dynabeads MyOne SILANE (Thermo Fisher Scientific). cDNA was then amplified with a C1000 Thermal Cycler programed at 98 °C for 3 min, 10 cycles of (98 °C for 15 s, 67 °C for 20 s, 72 °C for 1 min), 72 °C for 1 min, and hold at 4 °C. Subsequently, the amplified cDNA was fragmented, end-repaired, A-tailed, and index adaptor ligated, with cleanup in-between steps using SPRIselect Reagent Kit (Beckman Coulter). Post-ligation product was amplified with a T1000 Thermal Cycler programed at 98 °C for 45 s, 10 cycles of (98 °C for 20 s, 54 °C for 30 s, 72 °C for 20 s), 72 °C for 1 min, and hold at 4 °C. The sequencing-ready library was cleaned up with SPRIselect and sequenced by the Biomedical Sequencing Facility at CeMM using the Illumina HiSeq 3000/4000 platform and the 75 bp paired-end configuration.

### Assay for transposable-accessible chromatin with sequencing (ATAC-seq)

Chromatin accessibility mapping was performed using the ATAC-seq method as previously described^25,53^, with minor adaptations. In each experiment, a maximum of 50,000 sorted cells were pelleted by centrifuging for 5 min at 4 °C at 300 x g. After centrifugation, the pellet was carefully resuspended in the transposase reaction mix (12.5 μl 2xTD buffer, 2 μl TDE1 (Illumina), and 10.25 μl nuclease-free water, 0.25 μl 5% Digitonin (Sigma)) for 30 min at 37 °C. Following DNA purification with the MinElute kit eluting in 11 μl, 1 μl of the eluted DNA was used in a quantitative PCR reaction to estimate the optimum number of amplification cycles. Library amplification was followed by SPRI size selection to exclude fragments larger than 1,200 bp. DNA concentration was measured with a Qubit fluorometer (Life Technologies). Library amplification was performed using custom Nextera primers^25^. The libraries were sequenced by the Biomedical Sequencing Facility at CeMM using the Illumina HiSeq 3000/4000 platform and the 50 bp single-read configuration.

### Preprocessing and analysis of single-cell RNA-seq data

Preprocessing of the single-cell RNA-seq data was performed using Cell Ranger version 2.0.0 (10x Genomics). Raw sequencing files were demultiplexed using the Cell Ranger command ‘mkfastq’. Each sample was aligned to the human reference genome assembly ‘refdata-cellranger-GRCh38-1.2.0’ using the Cell Ranger command ‘count’, and all samples were aggregated using the Cell Ranger command ‘aggr’ without depth normalization. Raw expression data were then loaded into R version 3.4.0 and analyzed using the Seurat package version 2.0.1 with the parameters suggested by the developers^54^. Specifically, single-cell profiles with less than 200 detected genes (indicative of no cell in the droplet), more than 3,000 detected genes (indicative of cell duplicates), or more than 15% of UMIs stemming from mitochondrial genes were discarded. Read counts were normalized dividing by the total UMI count in each cell, multiplied by a factor of 10,000, and log transformed. The number of UMIs per cell and the percent of mitochondrial reads per cell were then regressed out using Seurat’s standard analysis pipeline.

### Dimensionality reduction and supervised analysis of gene expression

Principal component analysis, t-SNE analysis, hierarchical clustering, and differential expression analyses were carried out in R, using the respective functions of the Seurat package. t-SNE and cluster analyses were based on the first ten principal components. A negative binomial distribution test was used for differential analysis on genes expressed in at least 10% of cells in one group. Results were aggregated across patients by taking the mean for log fold changes and by Fisher’s method for p-values. Enrichment analyses were done using Enrichr API^55^ against the following databases: Transcription Factor PPIs, ENCODE, ChEA Consensus TFs from ChIP-X, NCI-Nature 2016, WikiPathways 2016, Human Gene Atlas, and Chromosome Location. Aggregate gene expression values for gene sets (signatures) were quantified as follows: Log-normalized transcript per 10^4^ UMI counts were scaled between 0 and 1. The values for all genes of a given set were then summed to obtain a raw value for each gene set and cell. To remove cell specific effects such as differences in UMI distributions due to sequencing depth, raw values were transformed to Z-scores using a distribution of raw values of 500 randomly picked gene sets of the same size. Differences in signatures between time points were assessed using ‘t.test’ in R. Results were aggregated across patients by taking the mean for log fold changes and by Fisher’s method for p-values. Multiple testing correction of differentially expressed genes, enriched terms, and differences in signatures was carried out using the Benjamini-Hochberg procedure as implemented by the ‘p.adjust’ function in R. The selected gene sets included 50 ‘hallmark signatures’ from MSigDB^56^, as well as ATF2, BATF, NFIC, NFKB1, RELA, RUNX3, and SPI target genes, and B cell signatures from Human Gene Atlas, NCI Nature 2016, and WikiPathways 2016, all obtained from Enrichr^55^. For data representation, we denoised the dataset with the deep count autoencoder (DCA) in Python^57^, using raw UMI counts as input and the ‘Zero-Inflated Negative Binomial’ model (which explained the relationship between mean expression and observed dropout rates significantly better than the ‘Negative Binomial’ model). The DCA-denoised data were then normalized per cell, log-transformed, and scaled. Dimensional reduction was performed by principal component analysis, and the resulting dimensions were used for neighbor graph construction followed by Uniform Manifold Approximation and Projection (UMAP) with Scanpy’s default parameters^58^.

### Preprocessing and analysis of ATAC-seq data

ATAC-seq reads were trimmed using Skewer^59^ and aligned to the GRCh37/hg19 assembly of the human genome using Bowtie2^60^ with the ‘-very-sensitive’ parameter. Duplicate reads were removed using the sambamba^61^ ‘mark-dup’ command, and reads with mapping quality >30 and alignment to the nuclear genome were kept. All downstream analyses were performed on these filtered reads. Peak calling was performed with MACS2^62^ using the ‘-nomodel’ and ‘-extsize 147’ parameters, and peaks overlapping blacklisted features as defined by the ENCODE project^63^ were discarded. We created a consensus region set by merging the called peaks from all samples across patients and cell types, and we quantified the accessibility of each region in each sample by counting the number of reads from the filtered BAM file that overlapped each region. To normalize the chromatin accessibility signal across samples, we first performed quantile normalization using the R implementation in the preprocessCore package (‘normalize.quantiles’ function). We then performed principal component analysis (scikit-learn, sklearn.decomposition.PCA implementation) on the normalized chromatin accessibility values of all chromatin-accessible regions across all samples. Upon inspection of the sample distribution along principal components, we noticed an association of several (but not all) samples from one processing batch with a specific principal component, while we did not observe any association of these samples with any known biological factor. To remove the effect of this latent variable while retaining variation from other (biological) sources, we performed principal component analysis on the matrix of raw counts on a per cell type basis (except myeloid cells, which contained no such samples) and removed the latent variable (first principal component) by subtracting the outer product of the transformed values of each sample in this component and the loadings of each regulatory element in the same component from the original matrix. We then again performed normalization of the corrected count matrix and component analysis jointly for all cell types as before.

### Time series modelling of chromatin accessibility dynamics

We modeled the temporal effect of ibrutinib in each cell type as a function of time by a latent process. To that end, we used the Python library GPy to fit Gaussian process regression models (GPy.models.GPRegression) on the log2 transformed sampling time on ibrutinib therapy (independent variable) and the normalized chromatin accessibility values for each regulatory element (dependent variable) for each cell type separately. We fitted a variable radial basis function (RBF) kernel as well as a constant kernel (both with an added noise kernel), and we compared the log-likelihood and standard deviation of the posterior probability of the two as previously described^64-66^. Dynamic regulatory elements were defined as those for which the survival function of the chi-square of the D statistic (twice the difference between the log-likelihood of the variable fit minus the log-likelihood of the constant fit) was smaller than 0.05 and the standard deviation of the posterior was higher than 0.05 (as described previously^66^). We then used the ‘mixture of hierarchical Gaussian process’ (MOHGP) method to cluster regulatory elements according to their temporal pattern. The MOHGP class from the GPclust library (GPclust.MOHGP) was fitted with the same data as before, this time with a Matern52 kernel (GPy.kern.Matern52) and an initial guess of four region clusters. Regions with posterior probability higher to 0.8 were selected as dynamic and included in the downstream analysis.

### Region set enrichment analysis

We performed region set enrichment analysis on the clusters of dynamic genomic regions using LOLA^30^ and its core database, which comprises transcription factor binding sites from ENCODE^63^, tissue-specific DNase hyper-sensitive sites^67^, the CODEX database^68^, UCSC Genome Browser annotation tracks^69^, the Cistrome database^70^, and data from the BLUEPRINT project^71^. Motif enrichment analysis was performed with the HOMER^72^ ‘findMotifsGenome’ command using the following parameters: ‘-mask -size 150 -length 8,10,12,14,16 -S 12’. Enrichment of genes associated with regulatory elements (annotated with the nearest transcription start site from Ensembl) was performed through the Enrichr API^55^ for the following databases of gene sets: BioCarta 2016, ChEA 2016, Drug Perturbations from GEO down, Drug Perturbations from GEO up, ENCODE and ChEA Consensus TFs from ChIP-X, ENCODE TF ChIP-seq 2015, ESCAPE, GO Biological Process 2017b, GO Molecular Function 2017b, KEGG 2016, NCI-Nature 2016, Reactome 2016, Single Gene Perturbations from GEO down, Single Gene Perturbations from GEO up, and WikiPathways 2016.

### Inference of global transcription factor activity

Global transcription factor accessibility was assessed by aggregating the normalized chromatin accessibility values of regulatory elements that overlap a consensus of regions (union of all sites) from ENCODE ChIP-seq peaks of the same factor across all cell types profiled. The mean accessibility of each sample in the sites overlapping binding sites of each factor was computed and subtracted by the mean accessibility of each sample across all measured regulatory elements. For visualization, we aggregated samples by cell type and sampling time point, displaying either the mean or a Z-score of chromatin accessibility. For all gene-level measures of chromatin accessibility, we used the mean of all regulatory elements associated with a gene, defined as the gene with the closest transcription start site as annotated by the RefSeq gene models for the hg19 genome assembly.

### Integrative analysis of ATAC-seq and single-cell RNA-seq data

To assess the agreement between the two analyses at the enrichment level, we performed enrichment analysis with Enrichr for genes differentially expressed across patients in the same cell type, and we compared the significance of terms for transcription factors in the ‘ENCODE TF ChIP-seq 2015’ gene set library with the significance of transcription factors enriched in the LOLA analysis for each ATAC-seq cluster. To identify a common transcriptional signature associated with ibrutinib treatment across cell types, we selected all genes that were differentially expressed with the same direction in at least 10 combinations of cell type and time point. These genes were split according to the direction of change with time and used for enrichment analysis with Enrichr as described above. The same genes were used to derive a score calculated as the mean expression of the upregulated genes over the mean expression of downregulated genes. An independent cohort of RNA-seq on bulk PBMCs from CLL patients^20^ was used to assess the reproducibility of the signature by observing the significance of the difference between scores upon ibrutinib treatment with a paired-samples t-test. To assess the performance of the score as a classifier, we generated a ROC curve by counting true positive and negative rates with a sliding score threshold, and we calculated the area under the curve with scikit-learn’s function ‘sklearn.metrics.auc’.

### Inference of DNA copy number variation from single-cell RNA-seq data

To infer DNA copy number profiles at the single-cell level, we started with DCA-denoised, normalized, and scaled single RNA-seq data of all cells. We removed per-cell differences by subtracting the median expression of each cell from all genes and per-gene differences by subtracting the median and dividing by the standard deviation. We then calculated a rolling mean of expression across genes ordered by their chromosomal position for each chromosome individually. To improve the representation of DNA copy number profiles, we centered the resulting matrix by subtracting the mean of all values in the matrix and applied smoothing by cubing the matrix values (which shrinks small changes relative to all cells) and multiplying them by 3 (which scales values back to usual copy number variation bounds). To discover clusters of genetically distinct cells within patients, we performed dimension reduction using principal component analysis on the smoothed matrix, computed a neighbor graph between cells, and fitted a UMAP manifold for the CLL cells of each sample (i.e. per patient and time point). This was overlaid with the response to ibrutinib of each single cell based on ibrutinib response signature described above. To assess global changes in genetic diversity within cells of a patient over time, we developed a global metric of genetic diversity based on inferred copy number profiles from single-cell RNA-seq data. We calculated pairwise Pearson correlation coefficients between all cells and used the square of the mean of this distribution as a measure of genetic diversity. To benchmark this approach, we first established simulated copy number profiles with the same dimensions are the inferred one but for varying total numbers of cells. We created two populations where we simulated gain or loss of chromosome 12 (log change: −1 or 1) whereas the remainder of the genome was Gaussian noise of mean zero and standard deviation 0.1. We assessed performance by mixing the two populations together in different ratios and computing Pearson correlation between the population fraction (ground truth) and the predicted global diversity. An additional benchmark was performed by taking advantage of natural, known mixtures of cell types in the data. For these data, the inferred change in genetic diversity is simply the difference between global diversity measures between time points of ibrutinib treatment for each patient.

### Prediction of sample collection and patient-specific response time from single-cell RNA-seq data

The time point of sample collection (day 0, 30, or 120/150) for each CLL single-cell transcriptome was predicted using the glmnet package in R with a multinomial response variable (for classification) and the ‘alpha’ parameter (lasso penalty) set to 1. Prediction performance was assessed by 3-fold cross-validation for each patient, where optimal ‘lambda’ parameters were obtained separately for each (outer) fold in a 5-fold inner cross-validation using the function cv.glmnet. Parameter ‘lambda’ for the final prediction across patients were obtained by 5-fold cross-validation on all data for each patient using cv.glmnet. Predictions were aggregated for each patient by taking the mean of dummy variables (1: early, 2: mid, 3: late) across the three other patients. Classification performance for support vector machines was assessed using the LiblineaR package in R. Classifiers were trained with ‘type’ parameter 0 and ‘cost’ parameters estimated by the heuristicC method on the training data, where cells were split ten times into 70% for training and 30% for testing. Quantitative prediction of the precise time (number of days) after the start of ibrutinib therapy was performed using the glmnet package in R with a Gaussian response variable (for regression) and the ‘alpha’ parameter (lasso penalty) set to 1. Prediction performance was assessed using the ‘lambda’ parameter that provided the highest R^2^ in the training data of each fold. The regularization parameter ‘lambda’ for the final prediction was obtained based on the mean squared error in a 3-fold cross-validation repeated five times on all data from each patient. Predictions were aggregated by taking the mean across three patients. For all Python analysis, we set the pseudo-random number generation seed state to 1142101101 in both the standard library ‘random’ and in ‘numpy’.

### Data availability

All data are available through the Supplementary Website (http://cll-timecourse.computational-epigenetics.org/). Single-cell RNA-seq and ATAC-seq data (sequencing reads, intensity values) have been deposited at NCBI GEO and are publicly available under accession number GSE111015.

### Code availability

The analysis source code underlying the final version of the paper will be provided on the Supplementary Website (http://cll-timecourse.computational-epigenetics.org/).

## Acknowledgements

We would like to thank all patients who have donated their samples for this study. We also thank the Biomedical Sequencing Facility at CeMM for assistance with next generation sequencing and all members of the Bock lab for their help and advice. C.S. was supported by a Feodor Lynen Fellowship of the Alexander von Humboldt Foundation. N.F. is supported by a fellowship from the European Molecular Biology Organization (EMBO ALTF 241-2017). T.K. is supported by a Lise-Meitner fellowship from the Austrian Science Fund (FWF M2403). D.A. and Cs.B. are supported by the K119950, KH17-126718, NVKP_16-1-2016-0004, and NVKP_16-1-2016-0005 grants of the Hungarian National Research, Development and Innovation Office, the Janos Bolyai research scholarship, and the LP95021 grant of the Hungarian Academy of Sciences. C.B. is supported by a New Frontiers Group award of the Austrian Academy of Sciences and by an ERC Starting Grant (European Union’s Horizon 2020 research and innovation programme, grant agreement n° 679146).

## Author contributions

A.F.R., T.K., D.A., C.S., and Ch.B. designed the study; S.T., M.R., Z.M., and Cs.B. provided samples and clinical data; T.K., F.Z., T.P., and C.S. performed the experiments with contributions from M.F., L.C.S., A.N., and D.A.; A.F.R., and N.F. analyzed the data with contributions from T.K. and Ch.B.; Ch.B. supervised the research. A.F.R., T.K., N.F., Z.M., Cs.B., D.A., C.S., and Ch.B. wrote the manuscript with contributions from all authors.

## Competing financial interests

The authors declare no competing financial interests.

## Supplementary figures

**Supplementary Figure 1:**
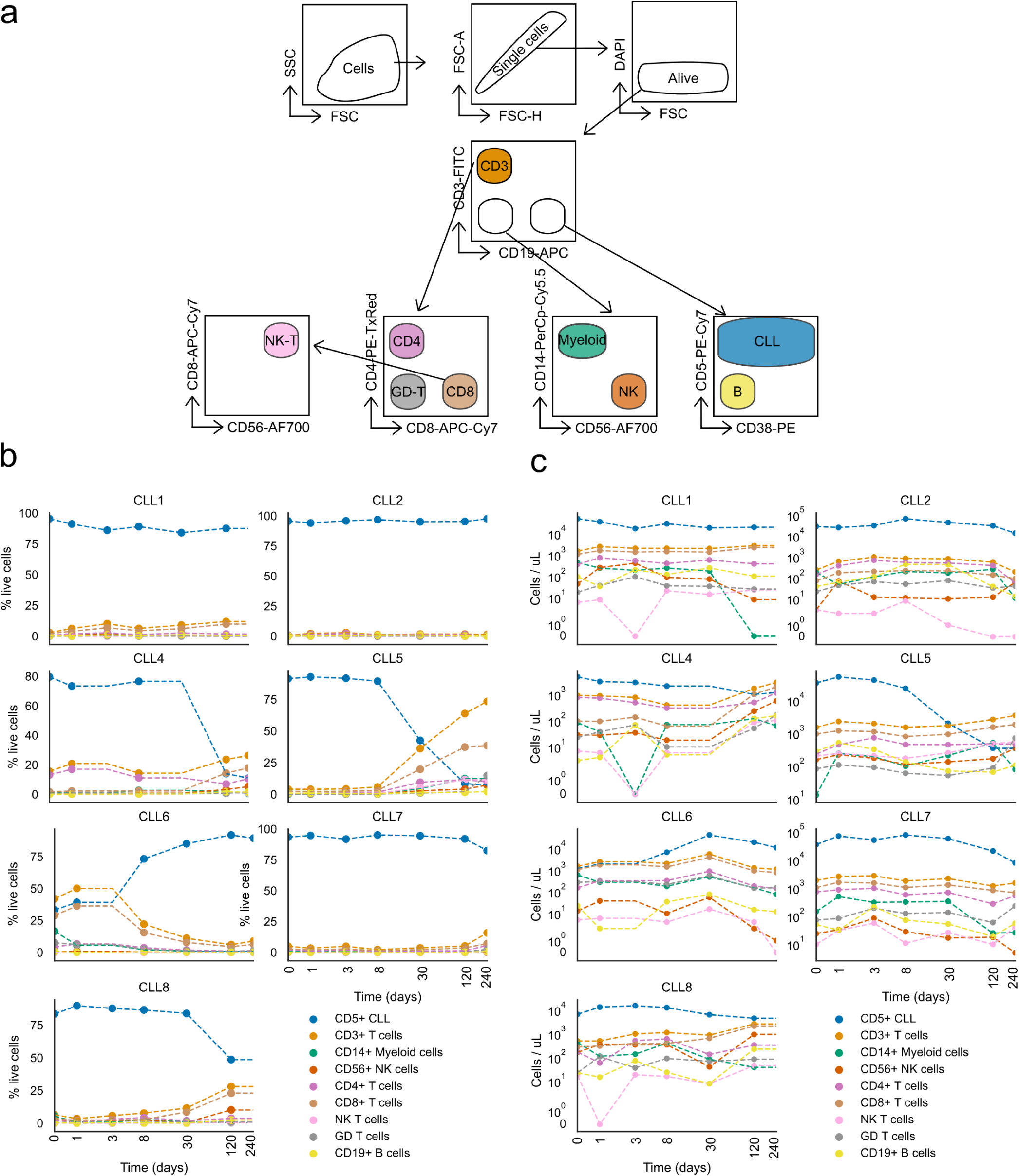
Temporal dynamics of cell composition in CLL patients upon ibrutinib treatment. **a)** Schematic representation of the FACS approach for purifying CLL cells and five non-malignant immune cell types from PBMCs of CLL patients. **b-c)** Flow cytometry based quantification of the relative (b) or absolute (c) abundance of CLL cells and several non-malignant immune cell types in patients undergoing ibrutinib therapy.

**Supplementary Figure 2:**
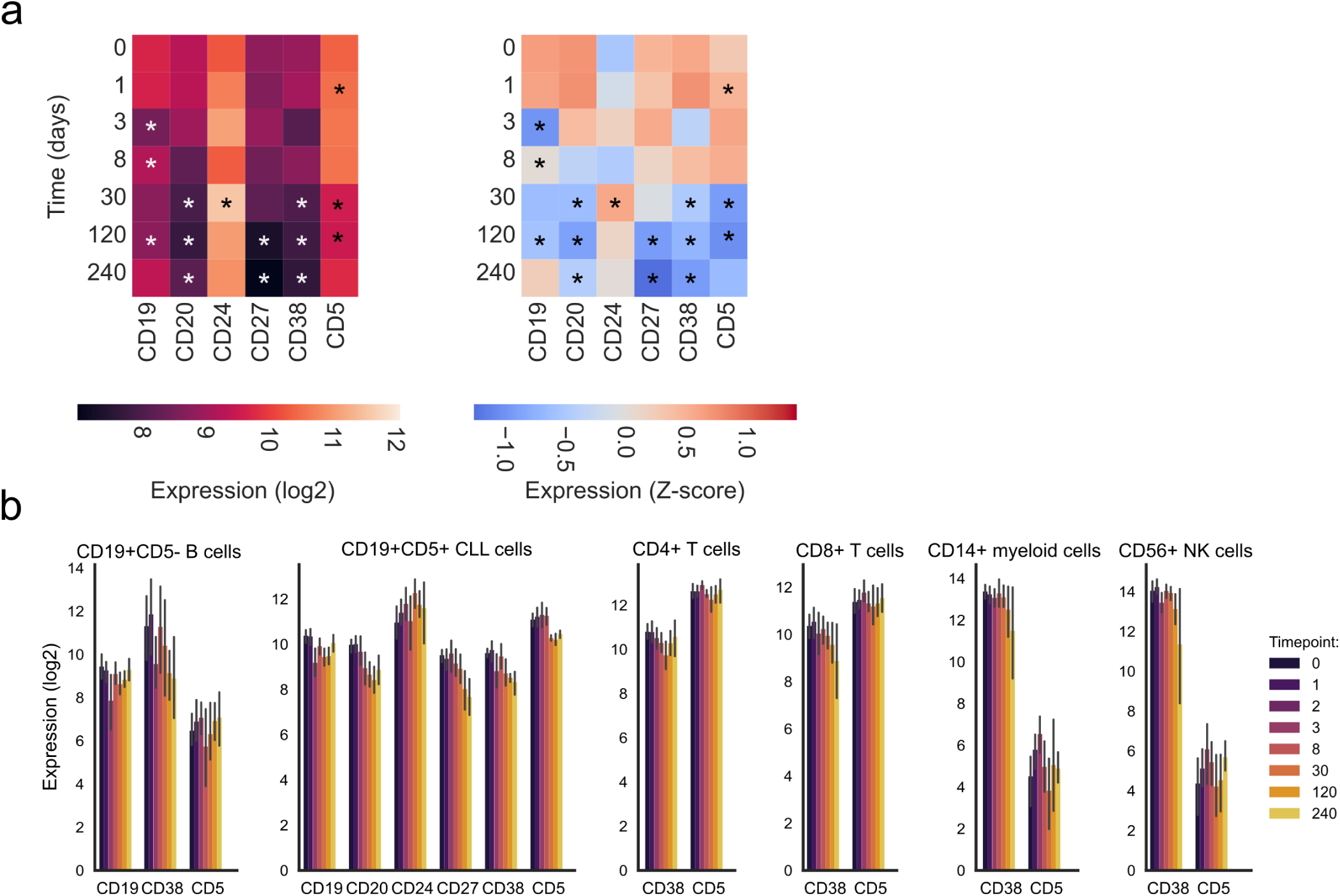
Temporal dynamics of T cell subsets and surface marker expression. **a)** Mean expression of surface marker proteins in CD19^+^CD5^+^ CLL cells during ibrutinib treatment. The left panel displays absolute (log scale) expression, while the right panel displays column-wise Z-transformed values. Stars mark significant changes compared to time 0 (paired t-test, p < 0.05). **b)** Expression of surface marker proteins in immune cell subsets of CLL patients as measured by flow cytometry.

**Supplementary Figure 3:**
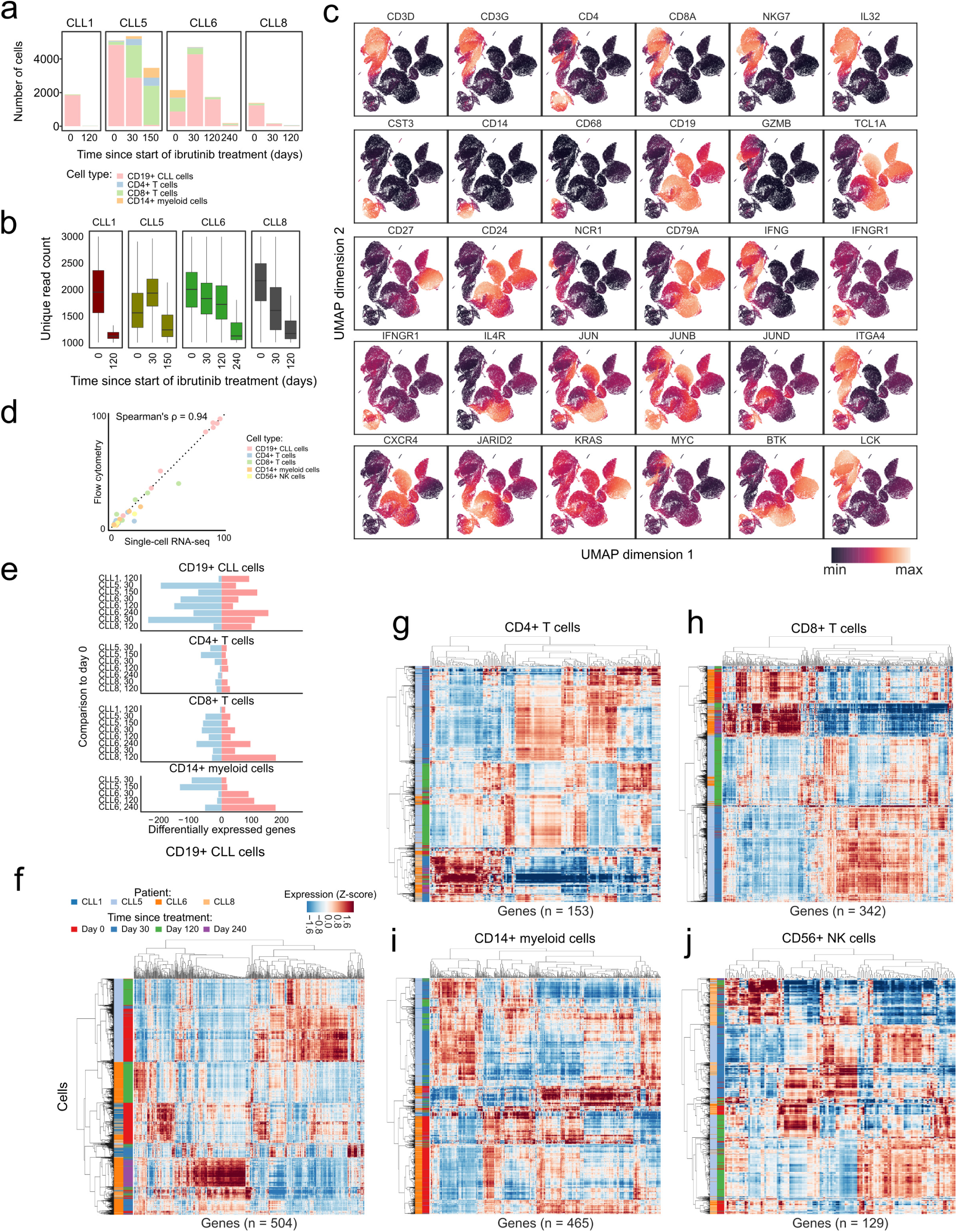
Single-cell RNA-seq profiling over the ibrutinib time course. **a)** Bar plot displaying the number of single-cell transcriptome profiles that passed quality control, shown separately for each patient, cell type, and time point. **b)** Box plot displaying the number of unique molecular identifiers (UMIs) detected per single cell, shown separately for each patient, cell type, and time point. **c)** Heatmap showing mean expression levels of the marker genes that were used to assign the single-cell transcriptomes to defined cell types. Values represent expression levels (normalized UMI counts) scaled from minimum to maximum in each row. **d**) Scatterplot comparing the fraction of cells of each type based on single-cell RNA-seq versus flow cytometry across all patients and time points. **e)** Number of differentially expressed genes for each cell type, patient, and time point. **f-j)** Heatmaps of differentially expressed genes over the ibrutinib time course separately for each cell type.

**Supplementary Figure 4:**
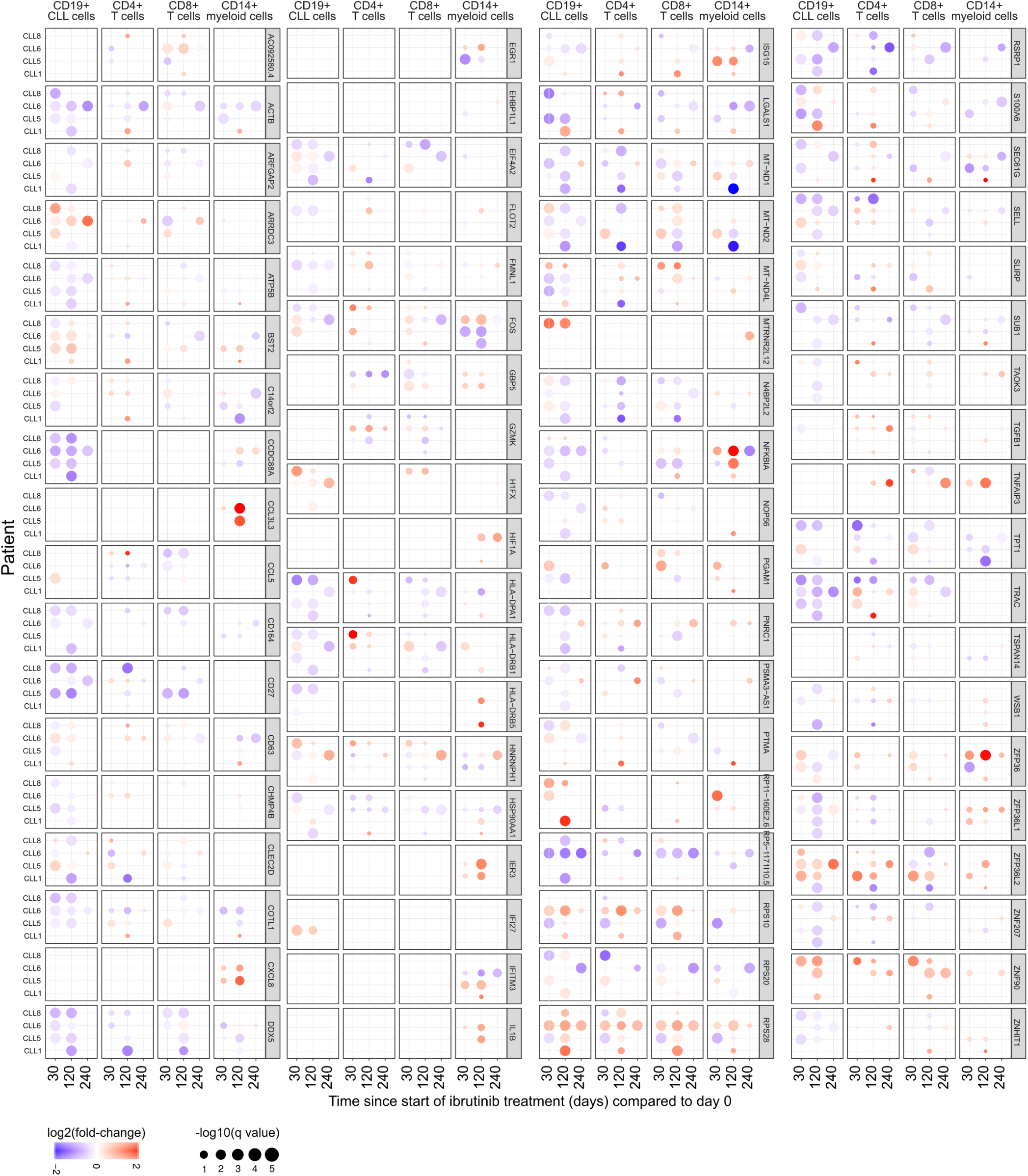
Transcriptional changes of differentially expressed genes upon ibrutinib treatment. Differential gene expression for those genes that were significantly differentially expressed (absolute log fold change >1 in more than two patients) over the course of ibrutinib therapy, shown separately for each cell type and comparing each sample to the pre-treatment (day 0) sample from the same patient.

**Supplementary Figure 5:**
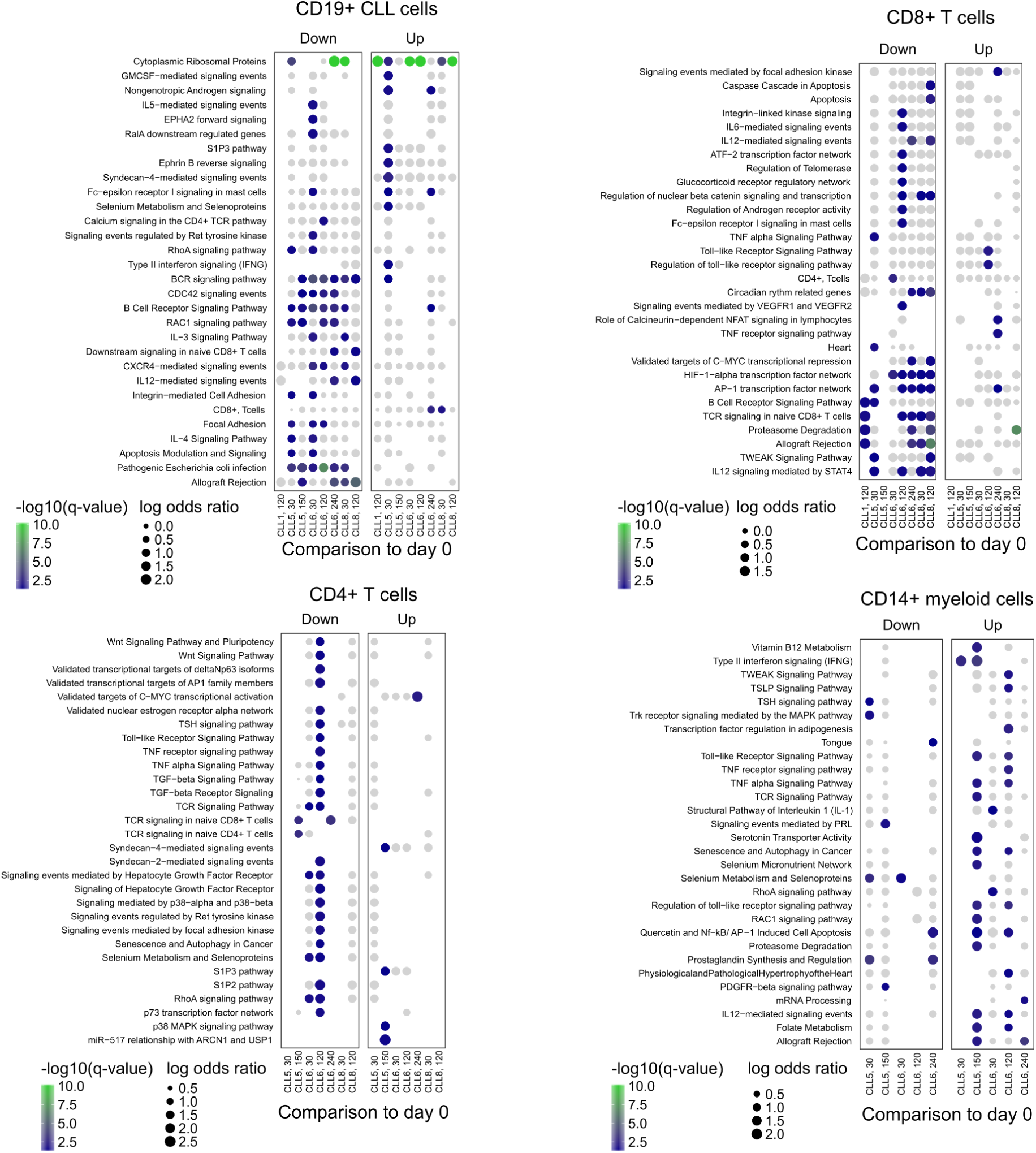
Gene set enrichments for single-cell RNA-seq over the ibrutinib time course. Enrichment of differentially expressed genes over the course of ibrutinib therapy for gene sets and biological processes involved in immune regulation, calculated separately for each cell type.

**Supplementary Figure 6:**
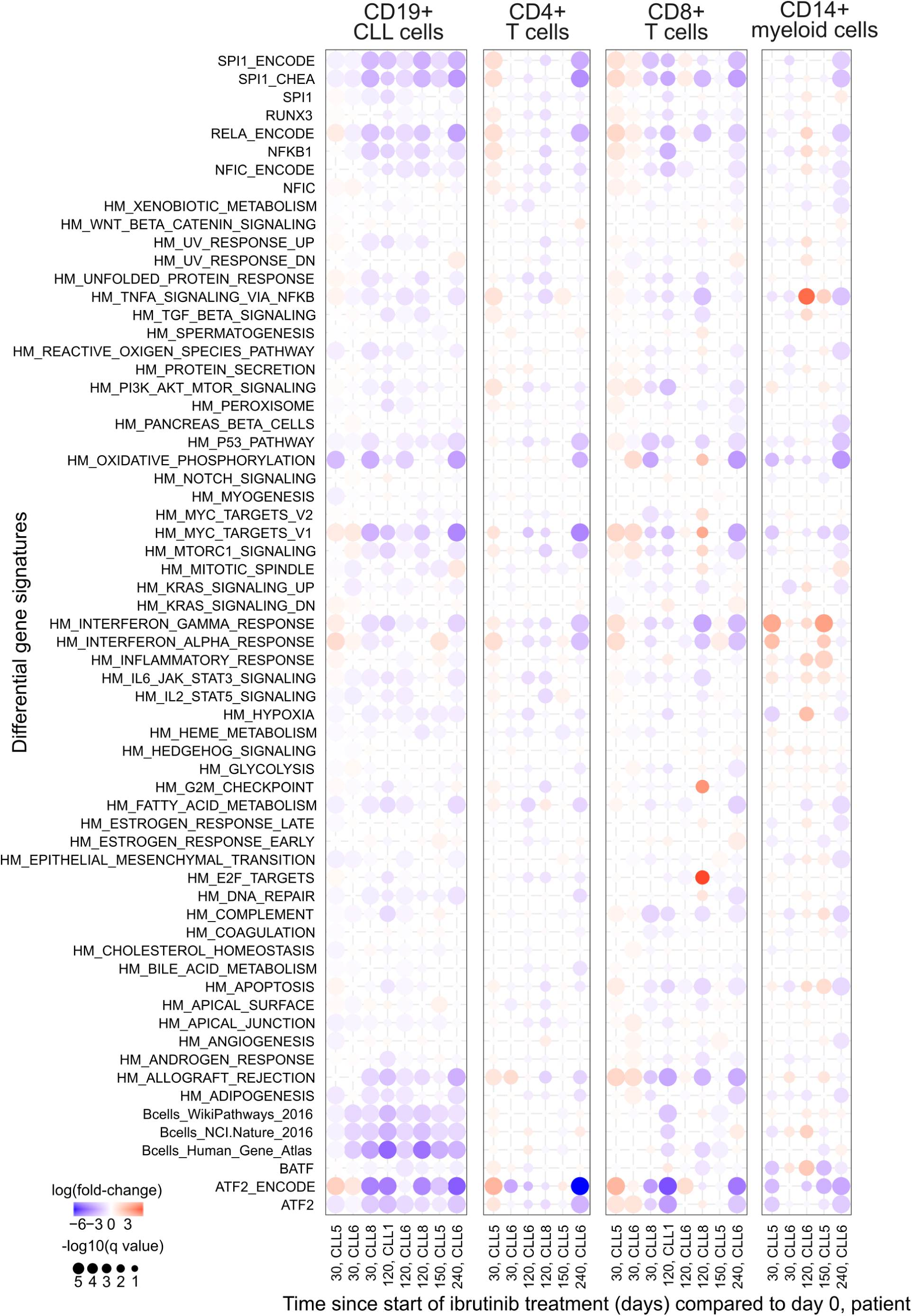
Transcriptional changes of selected gene signatures upon ibrutinib treatment. Aggregate gene expression of selected gene signatures plotted over the ibrutinib time course. The list of gene signatures includes the hallmark signatures from MSigDB^56^ (indicated by HM prefix) as well as sets of target genes for selected transcription factors (obtained from various sources).

**Supplementary Figure 7:**
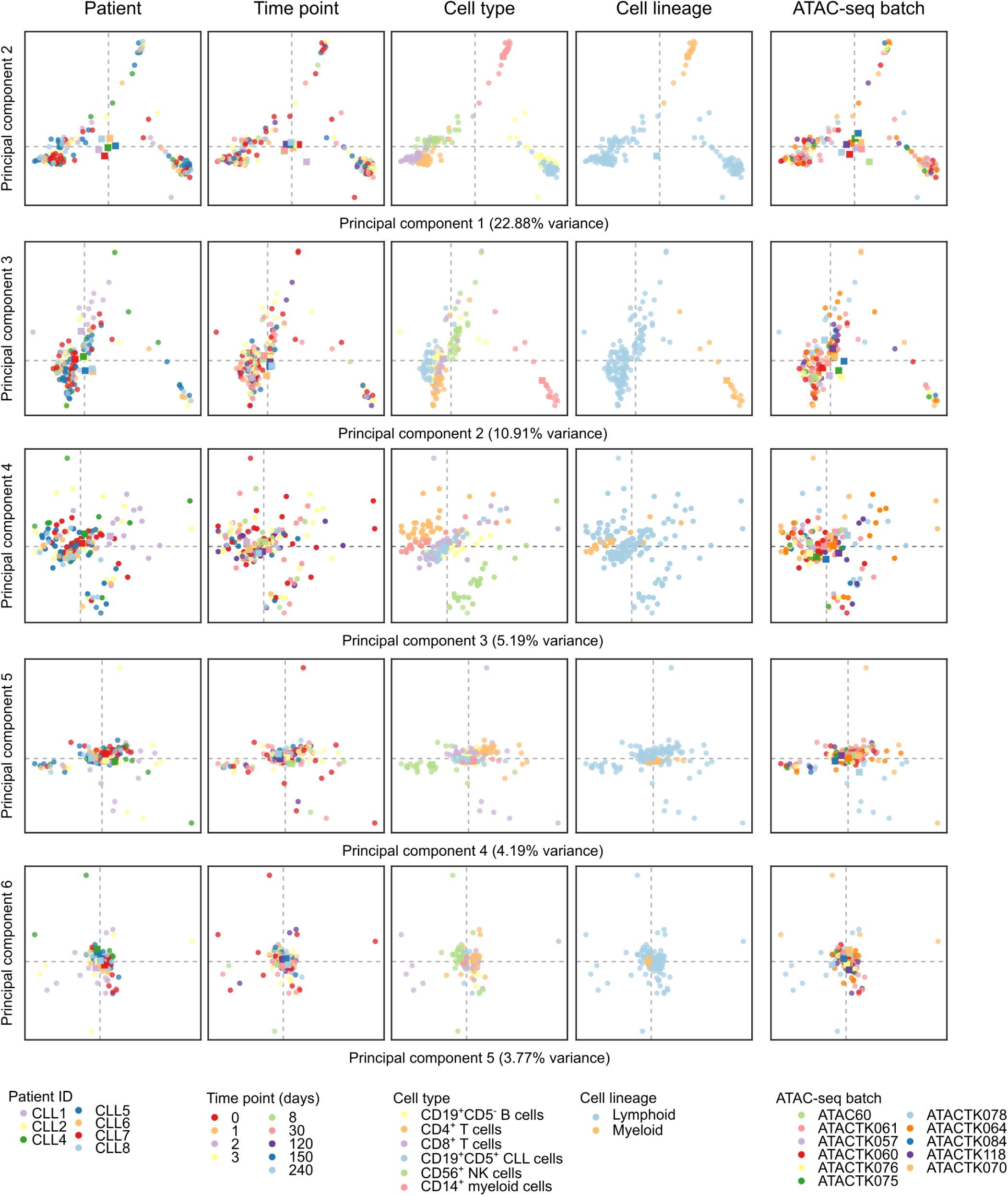
Unsupervised analysis of chromatin accessibility over the ibrutinib time course. Principal component analysis of all chromatin accessibility profiles, highlighting biological and technical annotations of potential relevance. Samples are shown as circles, color-coded according to the shown annotations, and the centroid for each annotation is shown as a color-coded square.

**Supplementary Figure 8:**
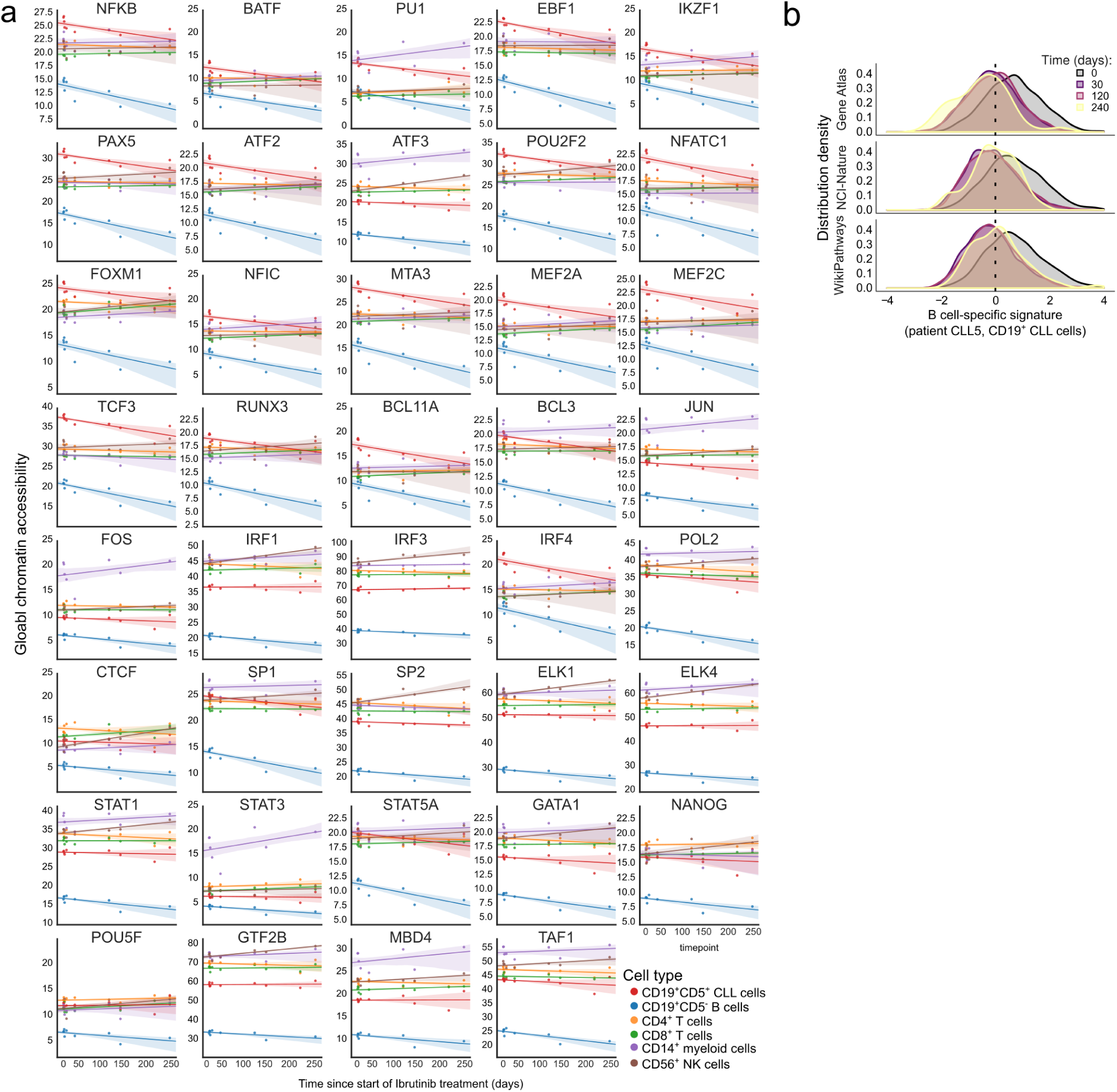
Changes in transcription regulation and cell state over the ibrutinib time course. **a)** Line plots showing mean chromatin accessibility of regulatory regions overlapping putative binding sites of the respective transcription factors (based on publicly available ChIP-seq data) for each cell type and time point. Colored areas indicate 95 percent confidence intervals calculated over 1,000 bootstrap runs. **b)** Gene expression histogram across CLL cells in one patient, demonstrating the decline of B cell-specific expression signature (three alternative signatures are shown) over the time course of ibrutinib treatment. For illustration, the patient with most time points in the single-cell RNA-seq analysis (CLL5) is displayed.

**Supplementary Figure 9:**
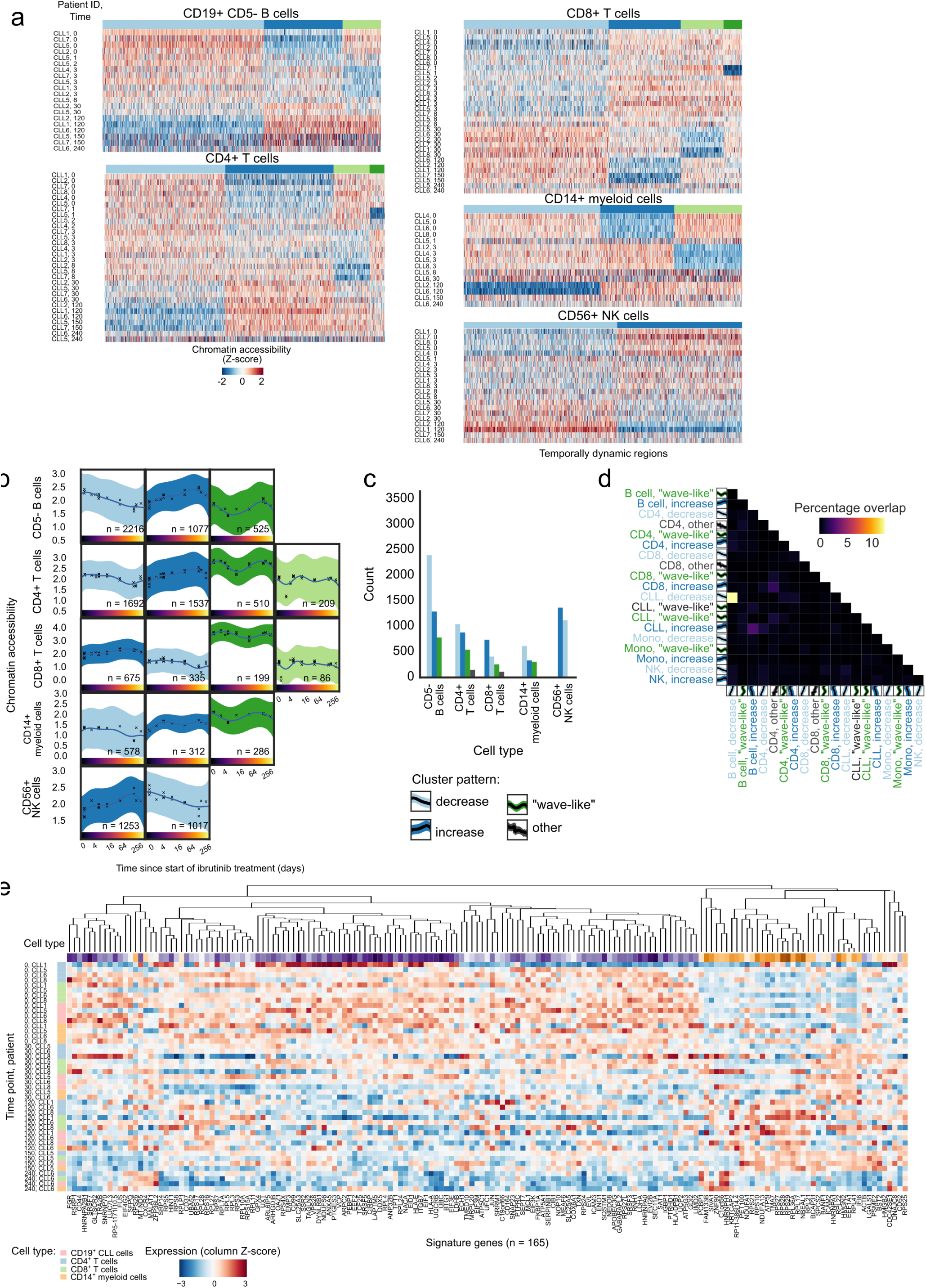
Cluster analysis of regulatory regions in non-malignant immune cell types. **a)** Heatmaps showing chromatin accessibility at dynamically changing regulatory regions for five FACS-purified non-malignant immune cell types collected over the ibrutinib time course. Values represent column Z-scores of normalized ATAC-seq signal strength. **b)** Mean chromatin accessibility across patients plotted over the ibrutinib time course for each cluster of dynamically changing regulatory regions in each cell type. Each cross represents a single sample from a single patient at a specific time point, and 95% confidence intervals are shown as colored shapes. **c)** Absolute number of dynamic regulatory regions for each cell type and cluster. **d)** Pairwise overlap of dynamic regulatory regions between cell types and clusters. **e)** Clustered heatmap showing patient-specific gene expression levels for the quiescence-like gene expression signature (**Figure 3e**), based on the single-cell RNA-seq data over the ibrutinib time course. Values represent column Z-scores of gene expression.

**Supplementary Figure 10:**
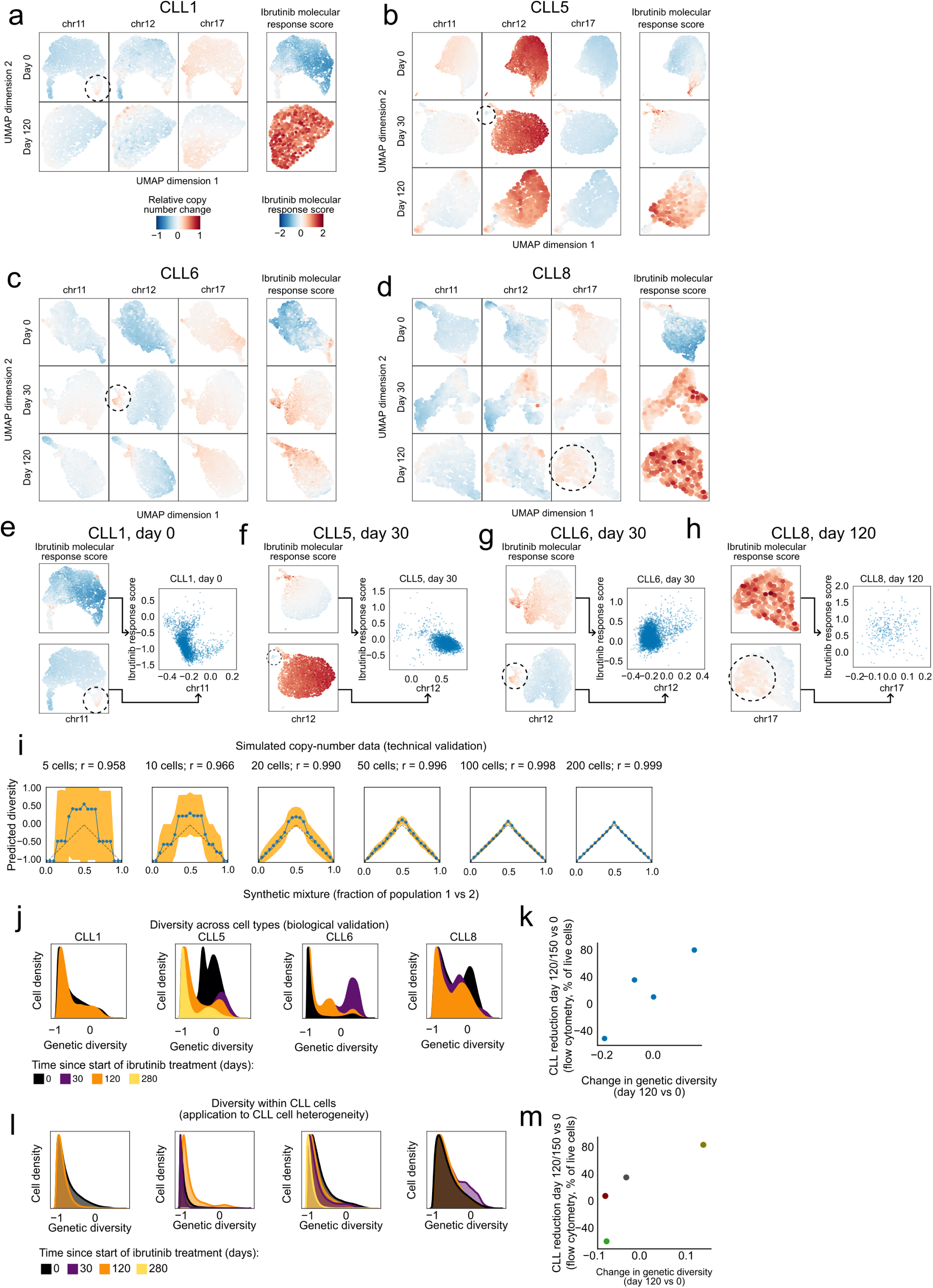
Analysis of copy number and genetic diversity over the ibrutinib time course. **a-d)** Two-dimensional similarity map (UMAP projection) based on DNA copy number profiles for single cells inferred from the single-cell RNA-seq data. These maps were calculated separately for each patient and time point. Color-coding indicates the relative copy number change for three chromosomal aberrations common in CLL (left) and for the ibrutinib molecular response score (right), i.e., the change in the CLL cell percentage on day 120/150 of ibrutinib treatment compared to day 0 as measured by flow cytometry. Genetically distinct subclones are highlighted by dashed circles. **e-h)** Scatterplots comparing selected subclonal copy number aberrations (x-axis and UMAP plots on the bottom left) with the ibrutinib molecular response score (y-axis and UMAP plots on the top left) across single cells in individual patients and time points. **i)** Accuracy of the computational approach for quantifying genetic diversity benchmarked on simulated copy number profiles that were combined at defined percentages (x-axis). Dashed lines indicate expected values (based on the simulation’s known ground truth), blue lines indicate inferred values, and yellow areas represents 95^th^ confidence intervals for the inferred values. Correlation coefficients quantify the overall agreement between expected and inferred values. **j)** Histograms showing the change in genetic diversity across all cells (i.e., CLL cells and immune cells) in the single-cell RNA-seq dataset. **k**) Scatterplot showing the correlation between the change in genetic diversity across all cells (x-axis) between time points and the cellular response to ibrutinib treatment (y-axis). **l**) Histograms showing the change in genetic diversity specifically for CLL cells. **m**) Scatterplot showing the correlation between changes in genetic diversity specifically in CLL cells (x-axis) and the cellular response to ibrutinib treatment (y-axis). Panel m is a reproduction of Figure 4b for consistency with panels j-k.

**Supplementary Figure 11:**
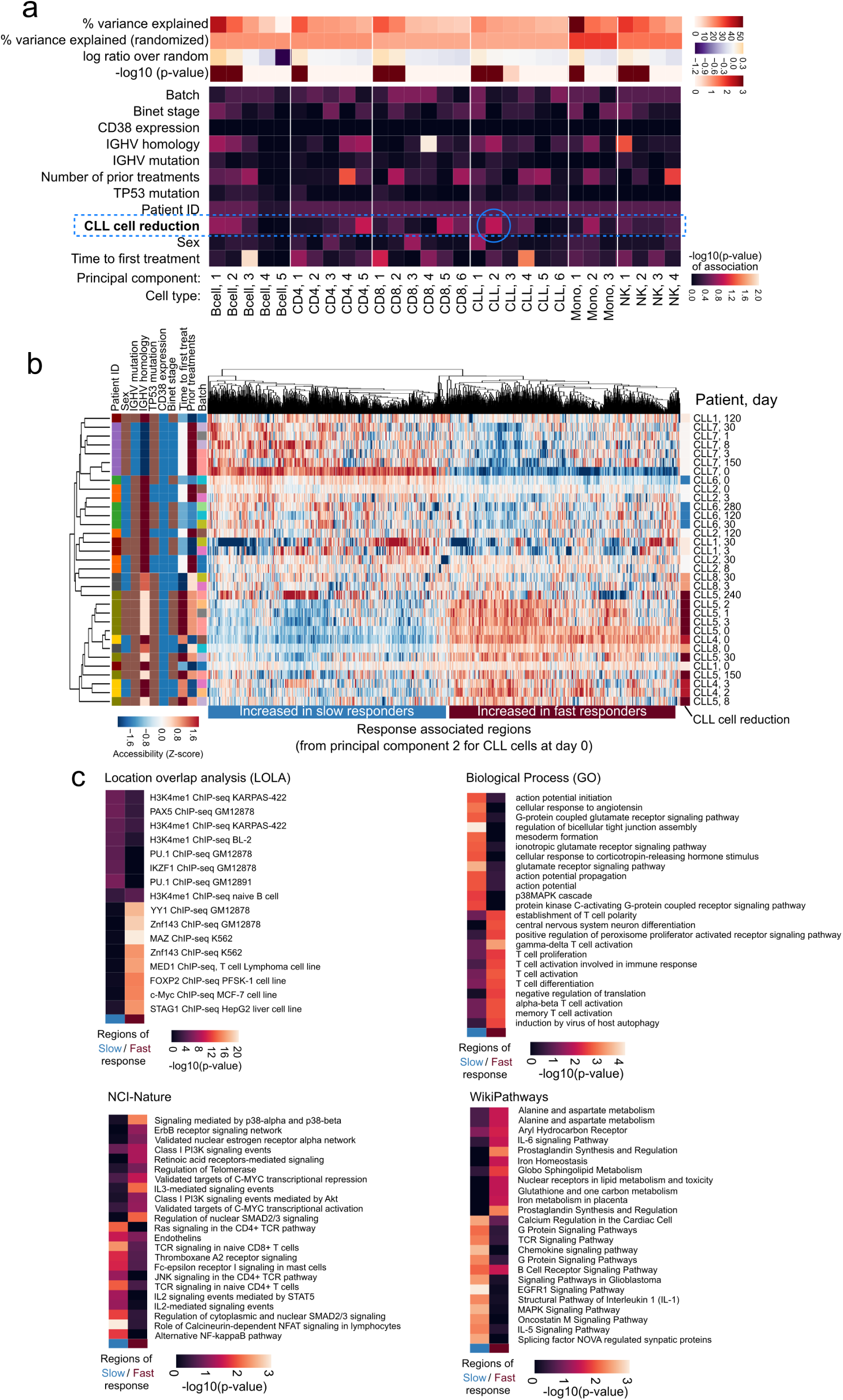
Analysis of chromatin profiles and their association with the response to ibrutinib. **(a)** Heatmaps showing the association of various clinical annotations with the principal components of the cell type specific chromatin accessibility profiles of different cell types prior to the start of ibrutinib treatment. The blue circle highlights the association between the second principal component for CLL cells and the change in the CLL cell percentage on day 120/150 of ibrutinib treatment compared to day 0, as measured by flow cytometry (y-axis). **(b)** Clustered heatmap showing patient-specific chromatin profiles for genomic regions associated with the second principal component (from panel a). **c)** Enrichment analysis for genomic regions associated with the second principal component, separately for regions associated with a slow versus a fast response to ibrutinib treatment.

**Supplementary Figure 12:**
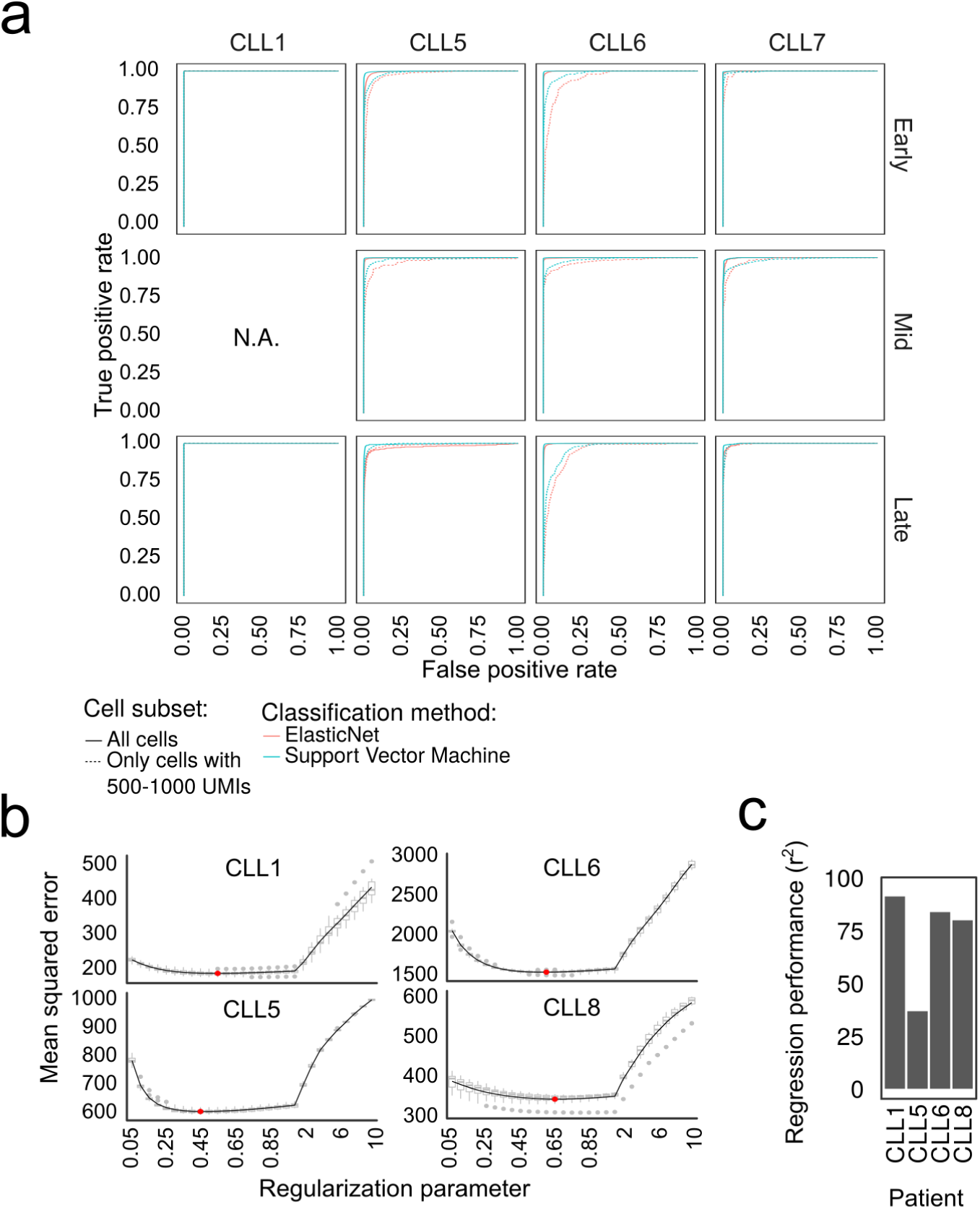
Prediction of the time point of sample collection from single-cell transcriptomes. **a)** ROC curves showing the cross-validated test set performance of classifiers predicting the time point of sample collection based on single-cell transcriptome profiles, using two different machine learning methods (logistic regression with elastic net regularization and support vector machines) and two different thresholds for single-cell RNA-seq data quality (all cells vs. only cells with 500 to 1,000 UMIs). **b)** Optimization of the regularization parameter (lambda) for predicting the time since the start of ibrutinib treatment using elastic net regularized linear regression. Red dots indicate the chosen parameter for each patient. **c)** Cross-validated test set performance of the regression models (coefficient of determination) for predicting the time since the start of ibrutinib treatment for each patient.

## Supplementary table legends

**Supplementary Table 1: Clinical annotation of the CLL patients included in the time course analysis**

**Supplementary Table 2: Cell type composition over the time course as measured by flow cytometry**

**Supplementary Table 3: Expression of cell surface marker proteins as measured by flow cytometry**

**Supplementary Table 4: Gene expression of individual cell types (single-cell RNA-seq) over the time course**

**Supplementary Table 5: Differentially expressed genes for individual cell types over the time course**

**Supplementary Table 6: Summary statistics for ATAC-seq chromatin mapping in CLL cells**

**Supplementary Table 7: Dynamic chromatin regions over the time course in CLL cells**

**Supplementary Table 8: Summary statistics for ATAC-seq chromatin mapping in immune cell types**

**Supplementary Table 9: Dynamic chromatin regions over the time course in immune cell type**

